# Plant actin networks are rapid mechano-adaptive scaffolds

**DOI:** 10.64898/2026.07.16.738884

**Authors:** Jasper Lamers, Anna Daamen, Ashlyn Pinto, Justin Tauber, Stefanie Gosselink, Cecilia Borassi, Uddalok Sen, Jasper van der Gucht, Joris Sprakel

## Abstract

The actin cytoskeleton is among the most highly conserved eukaryotic structures. In animal cells, actin plays a key role in mechanobiology, serving as a structural scaffold that shapes cellular mechanics and a mechanoresponsive network that reorganizes in response to mechanical cues. In plants, these roles are generally attributed to the cell wall and the microtubule cytoskeleton. This leaves the role of plant actin in mechanical responses largely unclear. Here, we ask whether a key actin feature that is essential for animal mechanobiology may also be conserved in plants. We use quantitative live-cell imaging and quantitative image analysis to show that plant actin networks are rapidly mechano-responsive. We find a quantitative correlation between the cellular distribution of cortical actin networks and cellular deformations induced by laser ablation. This process is rapid and completes within 2-4 minutes. We propose that this is an intrinsic physical property of crosslinked filamentous networks, as these findings can be reproduced in a physical model that lacks biological regulation. Moreover, actin networks also respond to different types of mechanical stresses: both osmotic treatment and squeezing lead to distinct changes in actin organization, distribution, and dynamics. Our findings show that plant actin networks form mechano-adaptive scaffolds that sense mechanical deformations and reorganize in response within just a few minutes. This gives rise to a picture of plants having a two-geared cytoskeletal mechano-response system in which microtubules mediate slow responses to subtle developmental stresses, whereas actin provides a rapid response to large and potentially damaging deformations.

## Introduction

The actin cytoskeleton is one of the most conserved structures in the eukaryotic domain (*1*). Not only is the actin protein itself highly conserved (*2*), but so is much of its core interactome (*3*), including actin nucleation, branching, and depolymerization factors, crosslinkers, and transport motor proteins. Given the high level of structural conservation, one may expect that some cellular functions of actin are also conserved. This is generally accepted to be true for actin’s role as a cellular transport highway and in mediating processes such as cytoplasmic streaming (*4*, *5*), while for other processes, such as actin contractility, it is clear that this is not conserved as, e.g., plants lack contractile myosins (*6*). In animals, actin also plays a central role in cellular mechanobiology (*7*). What the functional conservation of the mechanical functions of actin is beyond the animal domain is largely unclear. In animal cells, actin has two connected functions in cellular mechanics: i) it provides an elastic matrix that shapes the mechanical properties of the cell, and ii) acts as a mechanoresponsive structure that senses mechanical cues and reorganizes in response, for example, by accumulating at sites of high mechanical stress (*8–10*). Together, these properties form a robust system for rapid adaptation to mechanical challenges, whereby local stress-induced actin accumulation directly reinforces mechanically challenged regions (*10*, *11*). Consequently, actin occupies a central place in the study of animal mechanobiology.

The situation is markedly different in plants. Plant cell anatomy is dominated by the presence of a rigid carbohydrate cell wall. The inflationary pressure of cellular turgor acting against the elastic cell wall creates a balance of pushing and pulling forces—a so-called tensegrity balance—that provides the plant cell with its fundamental mechanical properties (*12*). Cell wall stiffnesses and turgor pressures can be as high as many MPa (*13*). By contrast, actin networks have stiffnesses many orders of magnitude softer, typically in the kPa range (*14–16*). As a result of this stiffness contrast, plant actin networks are generally thought not to play a significant structural mechanical role (*17*). Hence, relatively little attention has been paid to the possibility that plant actin networks act as mechanoresponsive structures capable of reading mechanical cues and organizing the cell interior in response (*18*). Instead, this function is typically attributed to the plant microtubule array, which reorganizes in response to subtle mechanical signals over the course of hours (*19*). Plant microtubules sense the mechanical field originating from the surrounding tissue (*17*, *19*) to steer the orientation of cell divisions and the anisotropic deposition of cellulose, thereby providing a mechanism for the mechanical control of tissue morphogenesis (*19–21*). Hence, in plant mechanobiology, microtubules have, to date, taken a central role.

In animal cells, which are soft and highly deformable, mechanical processes often involve large cellular deformations, and it is precisely in this context that the mechanical role of actin becomes most apparent. By contrast, cellular deformations in plant cells during developmental processes are typically much smaller and slower, as they are strongly attenuated by the rigid cell wall. This difference reinforces the view that actin plays little or no role in plant mechanobiology. However, several observations suggest that the prevailing view may be incomplete. Actin rapidly accumulates at sites of mechanical indentation caused either by pathogens or by abiotic indenters (*22*, *23*). The resulting actin patches appear to increase the mechanical barrier against pathogen penetration (*24*). Similarly, fungal pathogens, whose cells are themselves encased in rigid cell walls, assemble mechanoresponsive actin structures at sites of host contact, forming a structural scaffold that maintains tip-shape integrity to withstand the vast forces involved in host penetration (*25*). Evidence for a mechanical role of actin also exists in developmental contexts. Plant cell divisions are generally thought to align with the mechanical stress patterns acting upon cells, a phenomenon typically attributed to mechanosensing by cortical microtubules (*21*). However, distinct defects in mechanically instructed division orientations occur in the *act7* mutant of *Arabidopsis thaliana* (hereafter *Arabidopsis*) (*26*). This suggests that actin perturbation compromises either the perception of or the response to tissue-scale mechanical cues.

In this paper, we revisit the role of actin in plant cell mechanosensing and show that plant actin networks are highly and quantitatively mechano-adaptive under conditions of large cellular deformations. Using *Marchantia polymorpha* (hereafter *Marchantia*) gemmae, whose cells undergo large deformations following single-cell laser ablation, we demonstrate that actin networks reorganize as cells deform and local stress patterns change. We find a quantitative correlation between cortical actin accumulation and the degree of local cellular deformation. These responses are very rapid and complete within a few minutes. We also show that it is an intrinsic property of the actin network, independent of microtubules and driven by physics, as the results can be reproduced in a physical simulation model that lacks biological regulation. In Arabidopsis, laser ablation induces much smaller cellular deformations and correspondingly weaker actin responses. However, alternative mechanical perturbations, both squeezing and osmotic stress, show that *Arabidopsis* actin networks also alter their structure, distribution, and dynamics in response to mechanical signals. Together, our results support the view that plant actin networks form mechano-adaptive structural scaffolds that respond to large cellular deformations. These findings suggest that not only the actin protein and its core interactome but also key mechanical functions of the actin cytoskeleton have been conserved in the green lineage.

## Results

The structural and mechano-responsive nature of actin networks in animal cells manifest most prominently in scenarios where cells undergo large deformations. In plants, cell deformations are typically much slower and smaller as they are encased in a rigid elasto-plastic cell wall. To explore the mechanical function of plant actin networks, we began by searching for a model in which cellular deformations are relatively large when a single cell in a tissue is deflated by laser ablation. Here, we chose laser ablation as our method of choice, as it allows for the instantaneous induction of a localized mechanical imbalance within a multicellular tissue, immediate imaging after treatment for high temporal resolution of responses, and relatively high throughput for sufficient statistics. We find that laser ablation in various tissues in *Arabidopsis*, including the root (Fig. 1A) and hypocotyl epidermis (Fig. 1B), leaf pavement cells (Fig. 1C), and stomata (Fig. 1D), deformations of cells adjacent to an ablation site occur, but generally with low amplitudes (Fig. S1). While larger deformations were sometimes observed in Arabidopsis, the thin cell walls made it impossible to resolve from which cell the post-ablation signal originates (Fig. S2). By contrast, we find that ablating a single cell within the gemmae of *Marchantia* results in large bulging deformations with deflection amplitudes up to 5-7 μm (Fig. 1E). This suggests that while cell shape is largely hard-coded in the cell wall network in *Arabidopsis* and *Physcomitrium*, in *Marchantia* gemmae, it is determined to a much greater degree by the mechanical interactions between neighboring cells. This makes it a suitable starting point for our query. To explore actin responses in *Marchantia*, we generated two independent transgenic reporter lines for actin. One is based on the commonly used actin-binding peptide LifeAct (*pEF1α::LifeAct-mNeongreen*). Since LifeAct has been reported to disrupt actin structure and responses in some cases (*27*, *28*), we have generated a second independent line using the native *Marchantia* actin-binding protein MpADF3 (*pEF1α::ADF3-mScarlet3*). Both provide clear imaging of the actin networks in the cells of *Marchantia* gemmae (Fig. 1F-G), with no obvious differences in actin structure.

**Figure 1.**
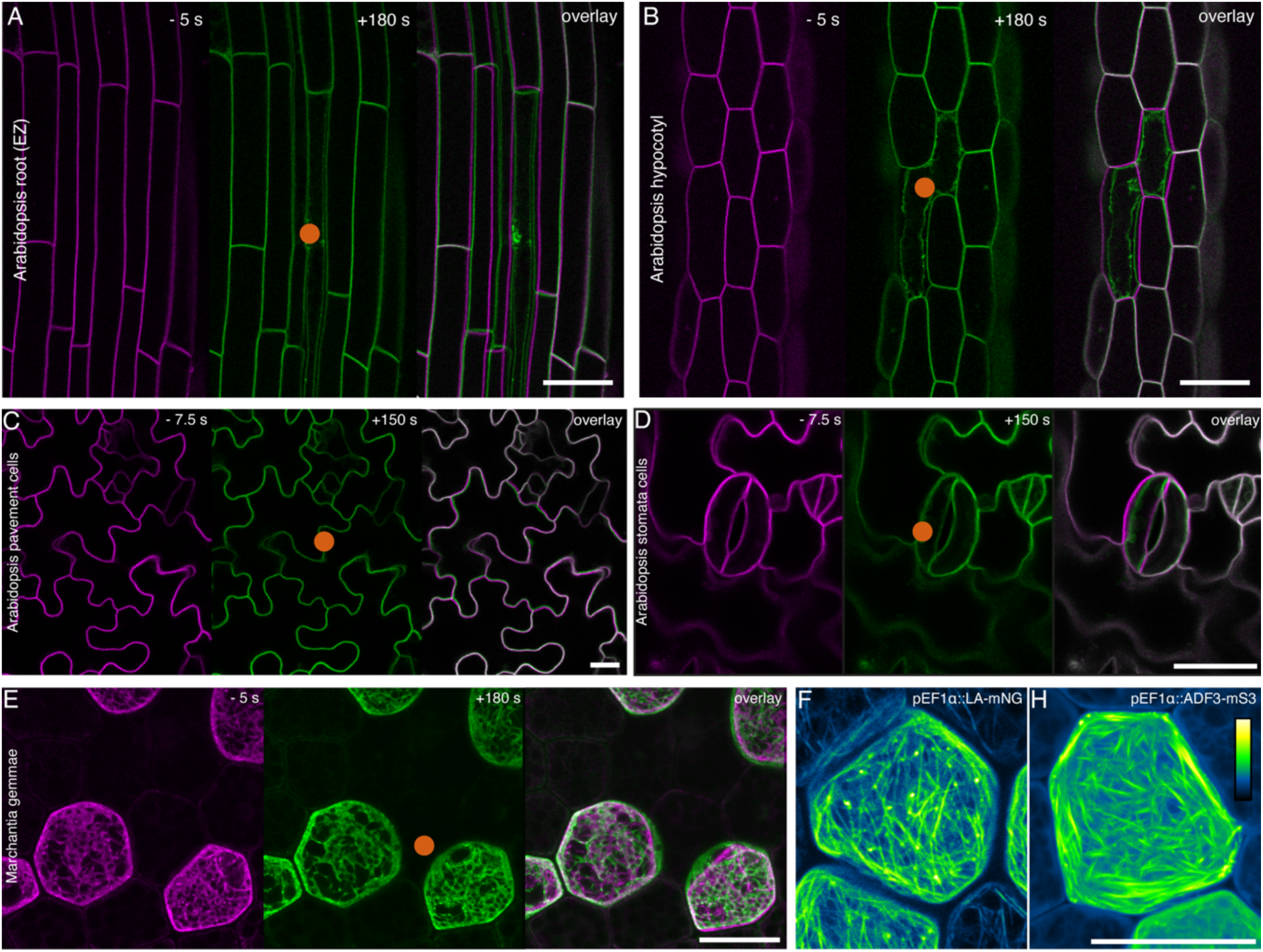
The amplitude of laser ablation-induced deformations is cell-type dependent. Various cell types of different organisms were ablated (A) elongation zone of an elongating root, (B) hypocotyl, (C) pavement or (D) stomata cells in Arabidopsis expressing p35::LTI6b-GFP, (E) gemmae cells of Marchantia expressing EF1α::LifeAct-mNeongreen. (F-G) Rhizoid precursor cells of Marchantia expressing EF1α::LifeAct-mNeongreen (F) and EF1α::ADF3-mScarlet3 (G). Deformation measurements are shown in Fig. S1. Red circles indicate the ablation point. Scale bar = 25 µm.

### Quantifying cellular deformations after laser ablation

Releasing turgor pressure from one cell via laser ablation creates a mechanical imbalance within the tissue. This causes cells surrounding the ablation site to undergo mechanical deformations to restore mechanical balance. We use these deformations as a readout of the cells’ mechanical response. To quantify the effect of the ablations, we developed a novel image analysis pipeline. This pipeline first binarizes the images in a time series (Fig. 2A) using Otsu binarization and then segments the cells into contours (Fig. 2B). Next, the centroid position of each enclosed cell contour is determined, from which a line is drawn to the cell outline to define the position of the cell contour (Fig. 2C). By comparing an image of the tissue before and during the ablation, the change in contour position can be quantified as the amount of cell contour displacement (Fig. 2D). This process is repeated for all pixels on the outer edge of each cell and performed for every cell in the image. In the same analysis, using an actin reporter, we can quantify its fluorescence intensity in the cell cortex. For this, we define a cortical mask within the cell contour (Fig. 2B); the contour edge is used to extract a region approximately 1 µm inward into the cell. All fluorescent signal within this strip around the cell contour is considered cortical actin. Also here, we defined the centroid position of each cell and drew a line to the cell edge to compute both the deformation and the change in fluorescence signal for each pixel along the cell contour, and for all rhizoid precursor cells neighboring the ablated cell. Data points were binned by their final deformation to enable performing statistics. Changes in actin intensity are computed from the ratio of intensities of the actin marker before, I(t<0) and after I(t) ablation as: 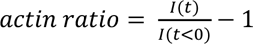. A value of 0 indicates that the actin intensity is unchanged, while negative values indicate that the actin signal is lowered because of the deformation, and positive values indicate that actin is recruited to that site. This yields clear results when considering a single cell (Fig. 2E-F) and can therefore be used for single-cell mechanical analysis. However, in the following, we averaged these data across multiple cells in independent experiments for statistical analysis. This script was validated on non-ablated samples, in which 5 pixels/timepoint were detected with >1 µm deformation versus 6241 without any deformation. There were no significant differences between these two groups (Fig. S3)

**Figure 2.**
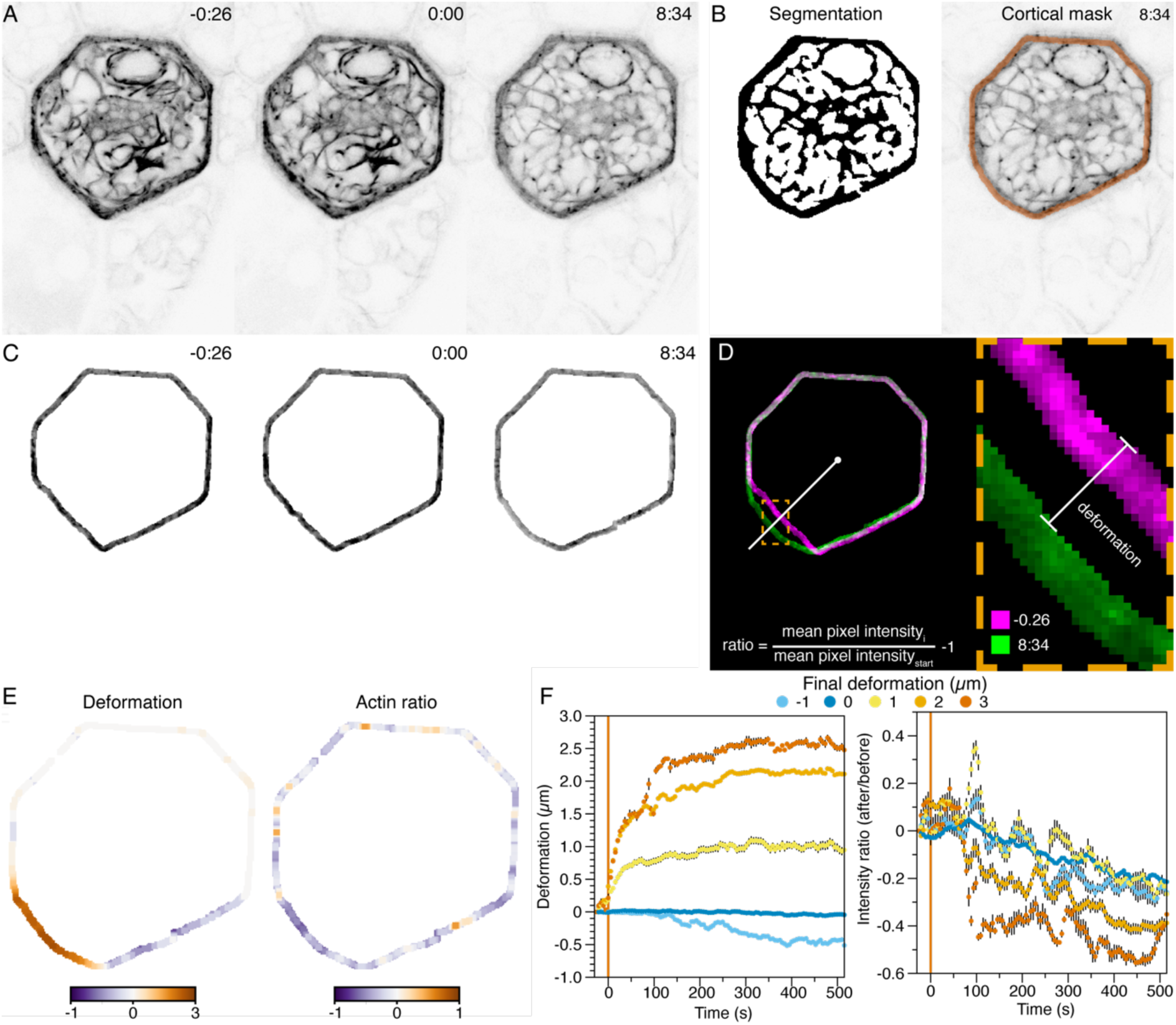
Deformation and actin intensity are calculated with an automated script. (A) Raw confocal images of the EF1α::LifeACT-mNeongreen at 3 different timepoints. (B) The segmented image after Otsu local thresholding and used to create a cortical mask (indicated in orange). For this example the +8:34 image of Panel A was used. (C) The cortical masks of the images in Panel A. (D) An overlay of the first and final timepoints of the series. For every pixel at the cell edge, a line is drawn to the center of the cell. The pixel values on this line are used to calculate the deformation and average intensity of the cortical fluorescent signal of a given length (1 µm as default). (E-F) The output of the deformation and fluorescent ratio intensity calculations. (E) Spatial maps resulted from the comparison between the first and last image of the timeseries or (F) on a temporal resolution when binned for the final deformation.

### Is Marchantia actin mechano-responsive?

We can now ask whether cortical actin networks in these *Marchantia* cells, analogous to their animal counterparts, are mechano-responsive and adapt their structure in response to mechanical deformations. To do so, we perform laser ablation experiments in a fluorescent actin reporter line (*pEF1α::LifeACT-mNeongreen*). Visual inspection of cells neighboring the ablation site clearly shows that the cell bulges outward into the gap left by the depressurized cell, while the actin simultaneously reorganizes and moves away from the bulging site (Fig. 3A).

**Figure 3.**
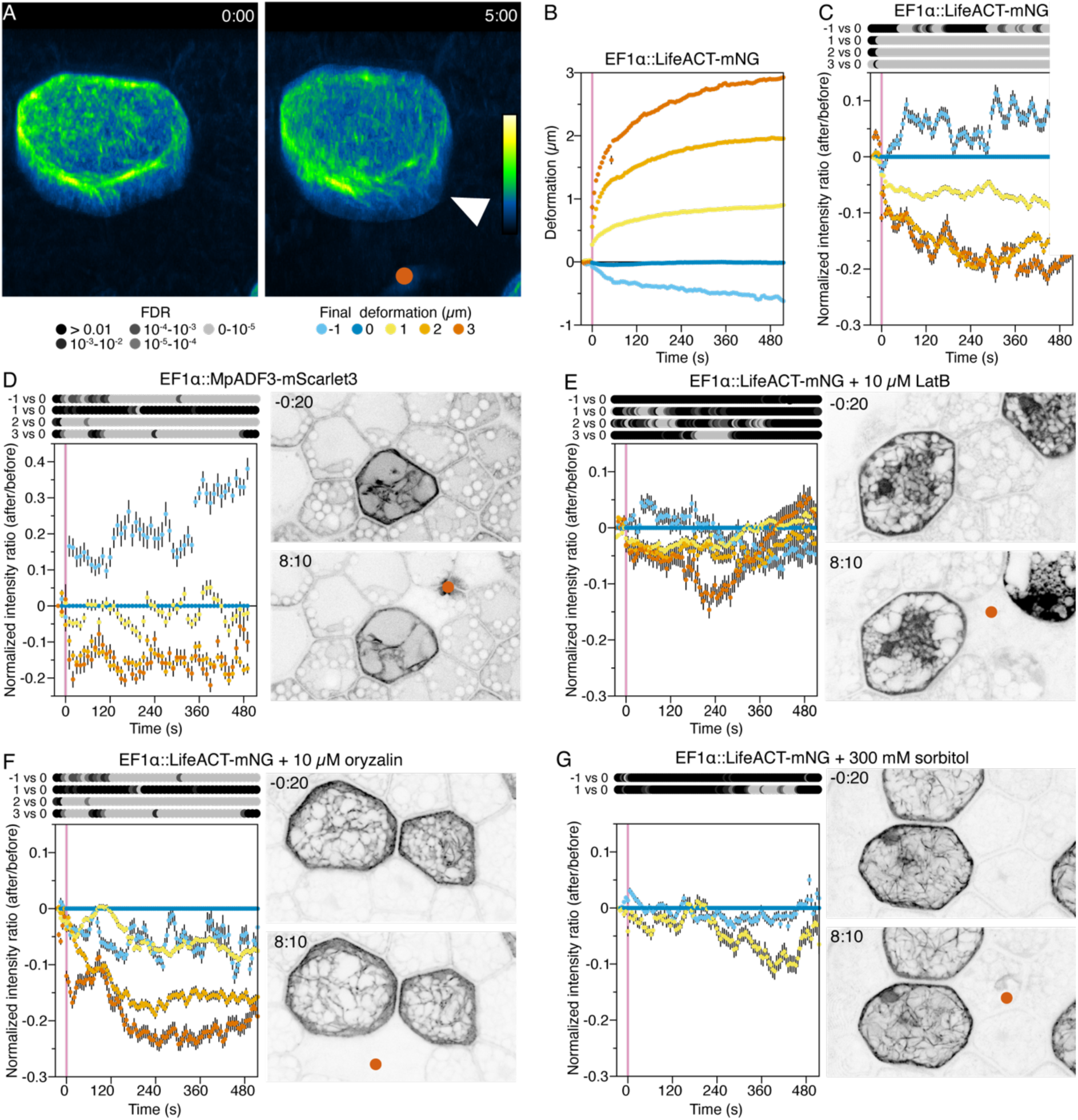
Cortical actin in Marchantia is mechanoresponsive. (A) 3D visualisation before and after a laser ablation of Marchantia expressing EF1α::LifeAct-mNeongreen from a 45° angle. (B) Deformations of the EF1α::LifeAct-mNeongreen line over time as binned on the final deformation of the timeseries. (C-G) Fluorescent intensity ratios over time, binned for the final deformation. (C) the EF1α::LifeAct-mNeongreen, (D) EF1α::ADF3-mScarlet3, (E) EF1α::LifeAct-mNeongreen treated with 10 µM LatB, (F) EF1α::LifeAct-mNeongreen treated with 10 µM oryzalin, (G) EF1α::LifeAct- mNeongreen treated with 300 mM sorbitol to limit deformations. Statistics were calculated with pairwise t tests per timepoint for every bin compared to 0 µm deformation. Multiple testing was corrected with Bonferroni’s method. Full details on sample sizes in Table S1. Non-normalized graphs are shown in Fig. S5. Red circle indicates the point of ablation and th1e arrowhead the bulging of the cell.

We mapped the dynamics of cellular deformation over time (Fig. 3B). Cells show a rapid initial deformation that occurs within a few seconds, followed by a slower creep-like deformation that takes minutes to reach the new equilibrium cell shape. This combination of a fast and a slow process is consistent with the mechanical response of a poroelastic material (*29*). Next, we quantified the actin polarisation ratio, as defined above. This shows that the actin rapidly moves away from the areas that show the most significant outward deformation; this response appears to be quantitative, i.e. the larger the deformation outward from the cell center, the larger the effect on the local actin concentration (Fig. 3C). These results were consisent and independent of the chosen width of the cortical region (Fig. S4). Since LifeAct, although often used, is known to cause artefacts in some cases (*27*, *28*), we performed the same experiments in an independent reporter line (*pEF1α::ADF3-mScarlet3*), which gave similar results (Fig. 3D). This suggests that plant actin networks are mechanoresponsive, and read mechanical cues from their environment and reorganize in response.

Cell puncture by laser ablation not only causes a mechanical imbalance in the tissue but can also induce thermal and light stress on the sample. In principle, cells in our field-of-view that do not deform are subject to these collateral processes serve as internal controls for these effects. However, to confirm that the actin reorganizations we observe are due to mechanosensing of the actin networks and not due to generic stress responses, such as cytoplasmic flows, we repeated the experiment in cells in which actin is disrupted with Latrunculin B, a toxin that inhibits actin polymerisation. We first visualized the effect of LatB and find that most of the filamentous f-actin becomes cytosolic g-actin, while only a few very thick actin bundles appear to resist the treatment (Fig. S6A). In LatB-treated cells, the mechano-responsivity of the actin is largely lost (Fig. 3E), indicating that the results we reported above are not due to generic ablation-stress responses or induced cytoplasmic flows.

Previously, a study on plant protoplasts in shaped microwells found that actin reorganization in response to geometric cues was dependent on the microtubule array (*18*). To explore whether what we observe is an intrinsic response of actin networks or also microtubule-dependent, we treated plants with oryzalin to depolymerize microtubule arrays. We confirmed by imaging in the microtubule reporter line *pMpEF1α::GFP-MpTUB1* (*30*) that the treatment effectively eradicates microtubule networks in gemmae (Fig. S6B). Cells without an intact microtubule network, however, still display a near-identical mechano-responsive behavior of cortical actin as in control experiments (Fig. 3F), indicating that these deformation-induced actin reorganizations are an intrinsic feature of the actin cytoskeleton itself. This is also consistent with the response kinetics of the two cytoskeletal systems. While the actin reorganizations we observe here occur within seconds to minutes, microtubule reorganizations in response to tissue stresses typically take hours (*19*).

Finally, to validate that the actin reorganizations are truly induced by the mechanical deformation of the cell, we manipulate the mechanical response of the tissue to laser ablation by reducing the effective turgor pressure in the cells by incubating the gemmae in a high osmolarity solution (300 mM sorbitol). In this case, cells indeed never deformed > 1 µm, and the actin response is also not observed (Fig. 3G). This, together with the LatB treatment, is another control indicating that the reduced actin in the most deformed areas is not an effect of the ablation itself but rather the result of a mechanical response system. We note that previous work on gametophore leaf tissues in *Physcomytrium* showed that actin accumulated near the ablation site, while there were almost no mechanical deformations there (*31*). Also, the time scale of this process was significantly slower. While we cannot explain this observation from our results, we note that this previous work lacked controls for laser-ablation-induced stresses or cytoplasmic flows, so it is possible that their observations reflect cytoplasmic expansion or ablation-induced stress responses at the wounding site rather than actin mechanoresponses.

Our data suggest that plant actin networks have the intrinsic capacity to read the cell’s mechanical state and reorganize in response. They do so quantitatively: the greater the deformation, the greater the response.

### A physical model for actin reorganizations

These results show that plant actin networks rapidly reorganize in response to changes in the mechanical field generated by the multicellular tissue. This raises the intriguing question of what the possible mechanism could be. Given the short timescale (tens of seconds) over which these reorganizations occur, this must result from a rapid-response system. We envision two, non-mutually exclusive, explanations: 1) actin is the client of a rapid mechanosensory regulatory system in the cells, e.g., involving the mechanical activation of kinases or actin-binding proteins (*8*, *32*), and/or 2) actin itself is mechanoresponsive without requiring biological regulation but based on physical principles (*33*). In animal cells, it is well established that both contribute to actin mechanosensing. However, most of the key regulatory components for option 1, such as integrins, associated mechanosensitive kinases, and catch bonds (*8*, *32*), have no known homologs in plants. While this does not rule out that plants have lineage-specific regulatory factors, we currently have limited handles to address this. Hence, we ask whether the observed actin reorganizations can be reproduced using a physical model alone, without biological regulation.

The model, described and illustrated in more detail in the SI, depicts a single rhizoid precursor cell adjacent to an ablation site. As a starting point, we extract an undeformed cell contour from a rhizoid precursor cell from experimental data prior to ablation. In this contour, we defined two distinct materials, each described as a quadrangulated mesh. The first mesh represents the cell wall, while the second represents the actin network. Actin is modeled in one of two limiting cases. In the whole-cell model, actin is represented as a continuum network that fills the entire cell. In the cortex model, we only define a cortical actin mesh and leave the cell interior empty. To simulate the ablation-induced deformation of the wall segment adjacent to the ablation site, the pressure in the cell is increased, while walls that remain connected to their neighbors are constrained from deforming. We then use a mechanochemical model that couples deformations in an elastic cell wall, due to pressure changes, mimicking what occurs in ablation experiments, to the molecular dynamics of actin filament and crosslinker binding.

We hypothesize that the actin relocalization upon cell deformation is due to force-induced cross-linker unbinding and/or actin depolymerization, both of which are established features of these networks (*34*, *35*). In essence, we assume that the biomolecular interactions that hold the actin filaments themselves and their crosslinkers together weaken as forces are applied to them. In the simulations, these mechanically induced dynamics are modeled using a kinetic reaction model that couples binding and unbinding to the mechanical strain at each site. Moreover, once unbound, crosslinkers can diffuse throughout the cell and bind at a new location. An increase in mechanical strain leads to more rapid dissociation of crosslinkers and subsequent diffusion, thereby reducing crosslinker concentration. In turn, reducing the crosslinker concentration lowers the stiffness of the actin network, which subsequently affects the mechanical strain. To parametrize the simulation, we take a stiffness ratio of approximately 1000 between cell walls (stiffness in the MPa range) (*13*) and actin networks (stiffness in the kPa range) (*14–16*), and create maximum deformations of 5 µm, which is consistent with the values obtained experimentally, as described above.

First, we note that this model, given the imposed constraints, recreates the bulging pattern of the rhizoid precursor cell when creating a pressure differential well (Fig. 4). We also observe that the induced deformation gives rise to a distinct, mechanically-triggered, relocalisation of the actin crosslinkers and actin network density in this model (color-coded patterns in Fig. 4). The actin density was measured in direct analogy to the experiment: we first divided the data into bins that quantify different levels of deformation, and within each bin, we averaged the actin intensity. As in the experiment, this was done for each time point in the simulation time series and by taking a cortical mask, hence only analyzing the cortical actin (even though in the whole cell model, a continuum elastic actin network is present across the whole cell). Interestingly, the model captures the experimentally observed trends very well. We observe a rapid decrease in actin in the zone adjacent to the ablation site, with a response amplitude quantitatively proportional to the local deformation (Fig. 4C,F). While the cortex-only model gives a qualitatively similar picture to the experiments, it shows significantly weaker coupling between deformation and actin relocalization. By contrast, the whole-cell model recapitulates the experimental data very well.

**Figure 4.**
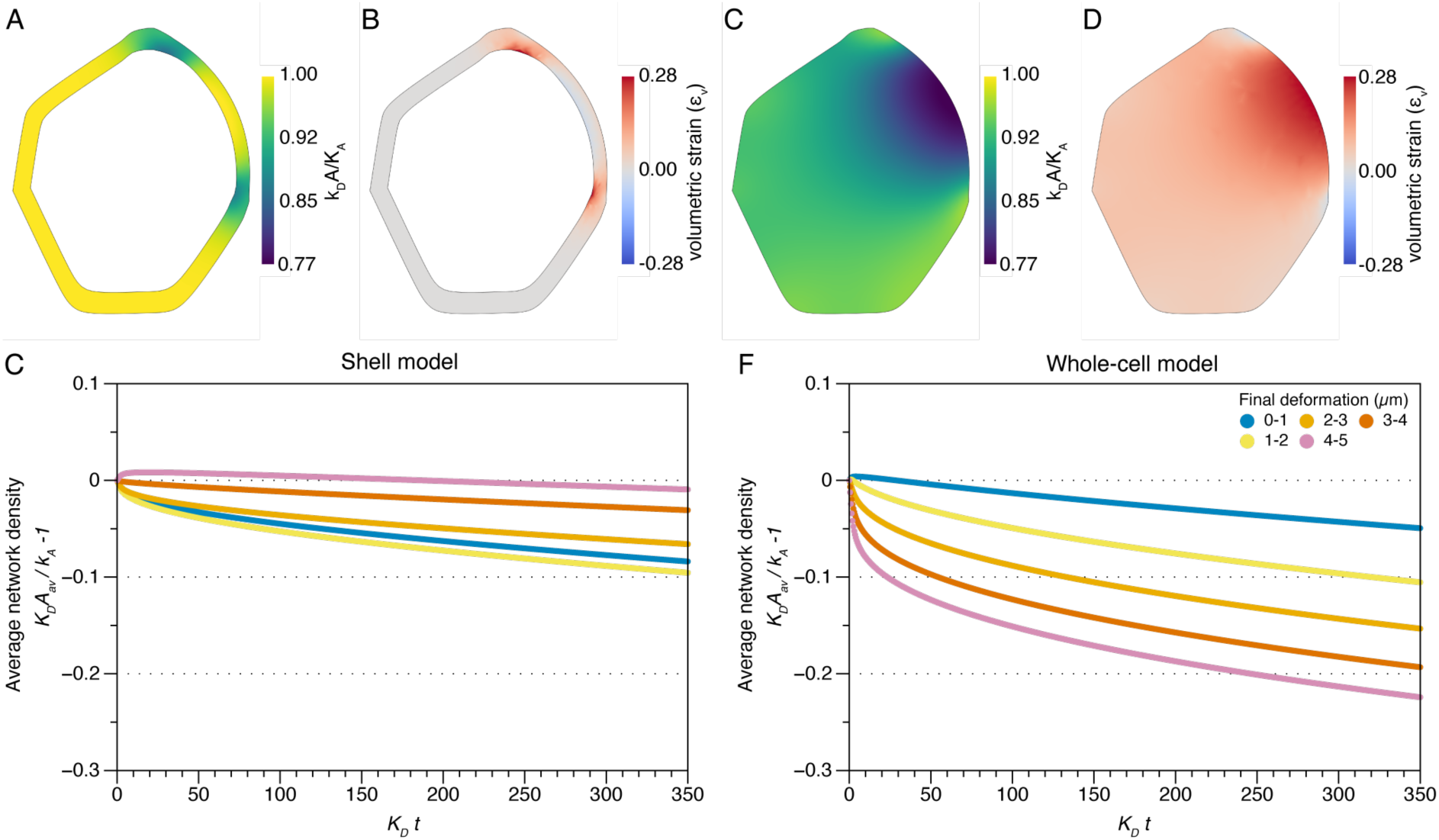
A physical model for actin reorganization upon cell deformation. (A-D) Heatmaps of the network density (k_D_A/k_A_) in the deformed cell for the shell and whole-cell model, respectively, and (B,E) represent heatmaps of the volumetric strain ε_v_ at the same timestep for the shell and whole-cell model, respectively. In the model we couple the volumetric strain to the actin network density and we observe that the network density decreases in regions of expansion. (C, F) Average network density (k_D_A/k_A_) measured over time for deformation bins of 1 µm each for the shell and whole-cell model respectively. Crosslinker concentration is relative to 1. A detailed explanation of the model and its quantities can be found in the SI.

These results lead to two interesting conclusions: 1) The experimentally observed actin relocalization can be quantitatively reproduced without invoking biological regulation, but by only describing cellular deformations and actin as a dynamically crosslinked network. As stated above, this does not exclude the possibility that biochemical regulation is involved, but it does indicate that a simple mechanochemical model is sufficient to explain the biological observations. Secondly, the model suggests that actin relocalization uses the entire cellular actin network as a mechanostat to sense local changes in tissue stresses, since a cortex-only model yielded much weaker couplings. This is important because we chose a model organism and cell type in which the vacuole occupies limited space, and hence, the actin networks forms a scaffold that spans much of the three-dimensional volume of the cell. This is very different for many other cell types in different plant species; for example, in the hypocotyl, leaf, and root epidermal cells in Fig. 1, the vacuole occupies the majority of the cell volume, leaving the actin networks restricted to a cortical cytosolic band around the cell. This may have implications for the cell type-specific sensitivity of actin networks to cellular deformations.

### Mechano-responsivity of Arabidopsis actin networks

So far, our findings have been obtained in the gemmae of the liverwort *Marchantia*. As an emerging model for land plant biology, *Marchantia* provides an attractive system for investigating the mechanical properties and functions of actin because of its relatively compact genome and limited gene family expansion. However, despite these advantages, its cytoskeletal organization and dynamics remain far less well characterized than those of the model angiosperm *Arabidopsis*. This raises the question of whether the rapid mechano-adaptive behavior of actin networks that we observe in *Marchantia* is conserved in *Arabidopsis*. As described above, *Arabidopsis* tissues show very limited deformation in response to laser ablation and exhibit less clear cortical actin accumulations. For this reason, we explored alternative approaches to explore the mechano-adaptivity of *Arabidopsis* actin networks.

We began by subjecting epidermal cells in the hypocotyl of an ABD2-mCherry (*36*) actin reporter line to squeezing, by confining the sample between two cover glasses, and to hyperosmotic shock. Both treatments place the cells under mechanical stress: one in a direct, directional manner (squeezing), while the other alters the tissue tensegrity balance not by removing a cell but by lowering the effective turgor pressure.

As a control for squeezing, we mounted hypocotyls in sample chambers with custom spacers, thicker than the specimen, to ensure they were freely suspended in the imaging chamber and not subject to squeezing forces at the glass-medium boundary. This is confirmed by 3D cross-sections of the hypocotyl surface in the spaced sample chamber (Fig. 5A) versus squeezed, and thus flattened, samples without spacer (Fig. 5B). We observed that the natural fluctuating dynamics of actin under control conditions, where actin undergoes relatively rapid, large-scale reorganizations over tens of seconds to minutes (Fig. 5C-D), were substantially slower and dampened in mechanically stimulated samples (Fig. 5E). We also note that these actin dynamics can be quite heterogeneous in the same sample between cells (Fig. 5D).

**Figure 5.**
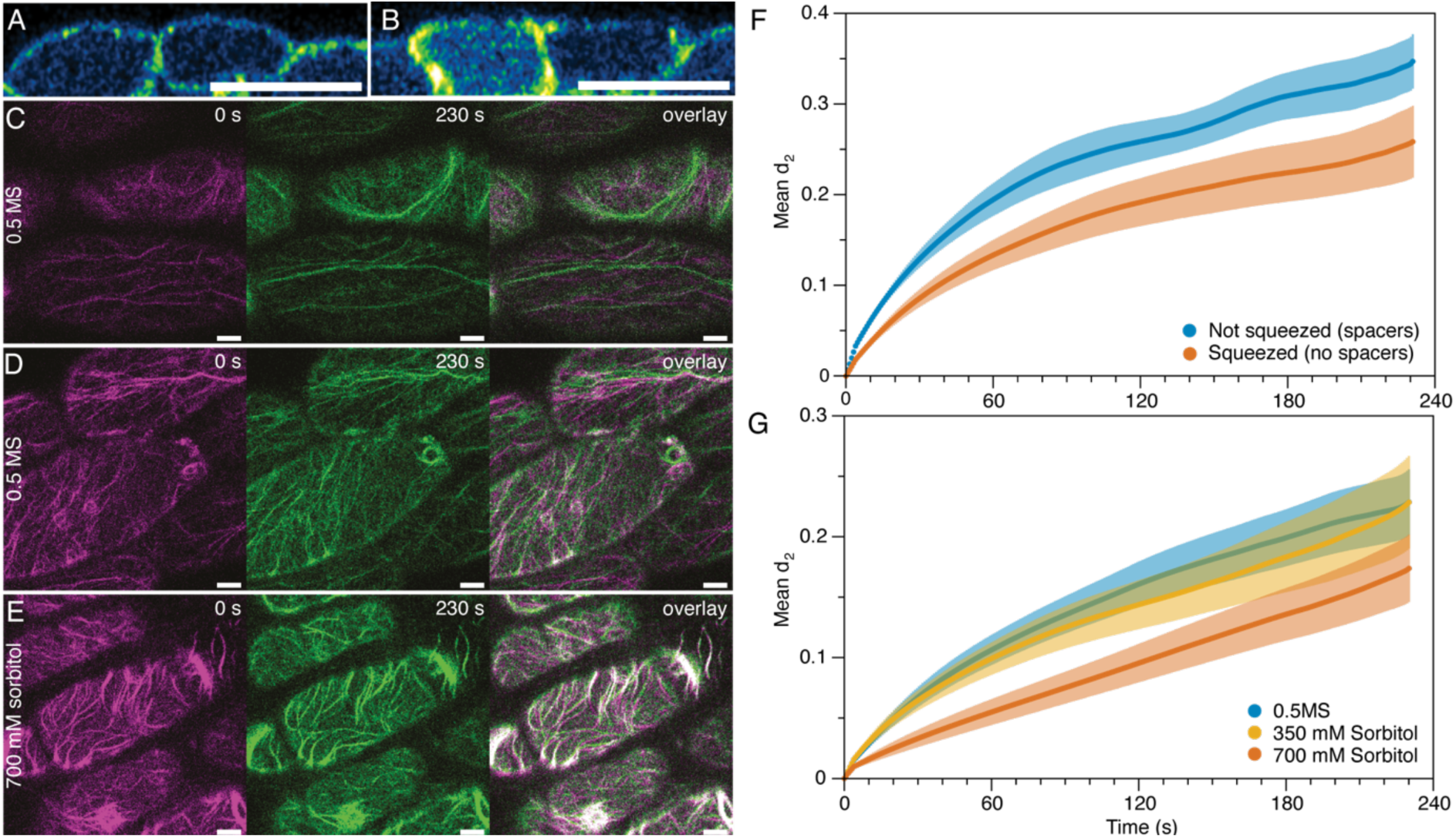
Mechanosensitive actin dynamics in Arabidopsis hypocotyl cells. A confocal cross-section with (A) and without (B) spacers. (C-E) First and last images of timeseries of actin in hypocotyl cells and their overlay: (C) 0.5 MS control, (D) 0.5 MS control that shows more heterogenous dynamics, (E) 700 mM sorbitol. (F-G) Actin dynamics quantified by the mean decorrelation over time for samples that were squeezed (F) or osmotically treated (G). Sample number not squeezed: n = 11, squeezed: n = 13, 0.5 MS: n = 14, 350 mM sorbitol: n = 16, 700 mM sorbitol: n = 12. Scalebar = 25 µm.

We quantified actin dynamics using an image-correlation method derived from Laser Speckle Imaging. In this analysis, two images separated by a time interval Δt, are compared by computing the extent of image decorrelation, which is quantified using a correlation function: the intensity structure function *d*_2_. At short time intervals, where actin has moved little, two images are very similar, and the decorrelation value is close to zero. At larger time intervals, where actin has had more time to move, decorrelation increases. The rate and extent of decorrelation over time reflect the actin dynamics. For control conditions, in the spaced sample chamber, the decorrelation grows rapidly over the course of one to two minutes (Fig. 5F), reflecting the rapid intrinsic fluctuations of actin turnover in the filaments and bundles in the cell. We find that all mechanical treatments decelerate actin dynamics, as reflected by a reduced decorrelation in our analysis (Fig. 5F-G), consistent with our visual observations. Squeezing yields a reduction in actin dynamics compared to the control samples mounted with a spacer (Fig. 5F). Also, for the osmotic treatment, we observe a dose-dependent effect, with 700 mM sorbitol yielding greater attenuation of the dynamics than 350 mM (Fig 5G). The sample-to-sample variation across different samples within the same treatment or control group is relatively large, as described above (Fig. 5D), and is shown in Fig. 5F-G as an error cloud indicating the standard error between replicates. Yet, we observe that the treatment-induced changes in dynamics for the squeezing and the highest osmotic stress exceed the control sample’s error cloud, i.e., the effect of this treatment exceeds the intrinsic variation within a sample group. Based on this, we evaluate these effects as significant reflections of mechanics-induced changes in actin dynamics and a first indication that actin networks in *Arabidopsis* are also mechano-adaptive. Interestingly, these data also suggest that both squeezing and osmotic stress have similar effects on actin dynamics.

In these experiments, we also observed, as shown in Fig. 5D, a correlation between the appearance of the actin network and its dynamics. Cells with larger and thicker bundles appear to be substantially less dynamic than those that primarily show finer actin filaments. Hence, this made us wonder whether the reductions in actin dynamics we observe here result from mechanically induced changes in the spatial organization of actin, as we also observed and quantified in *Marchantia* (Fig. 3). To explore this, we performed more detailed imaging experiments using the same ABD2-mCherry reporter line.

To study how osmotic stress impacts actin organization during large scale cell deformation, we plasmolysed hypocotyl cells which leads to severe cellular deformations as the protoplast recedes from the cell wall over the course of tens of minutes. In the control samples (mounted in isotonic 0.5 MS medium), actin is observed to fluctuate, as also described above (Fig. 5), but no large scale rearrangements are observed on the time scale of this experiment (Fig. 6A). By contrast, when deformations of the membrane are induced by plasmolysis, actin is rapidly enriched at the sites of largest deformation (Fig. 6B, some events indicated with arrows). This is consistent with previous reports (*37*). Also here, similar to our observations for actin dynamics, we observe large cell-to-cell variations within the same sample, although the general trend is very clear; large inward cellular deformations lead to significant actin accumulation.

**Figure 6.**
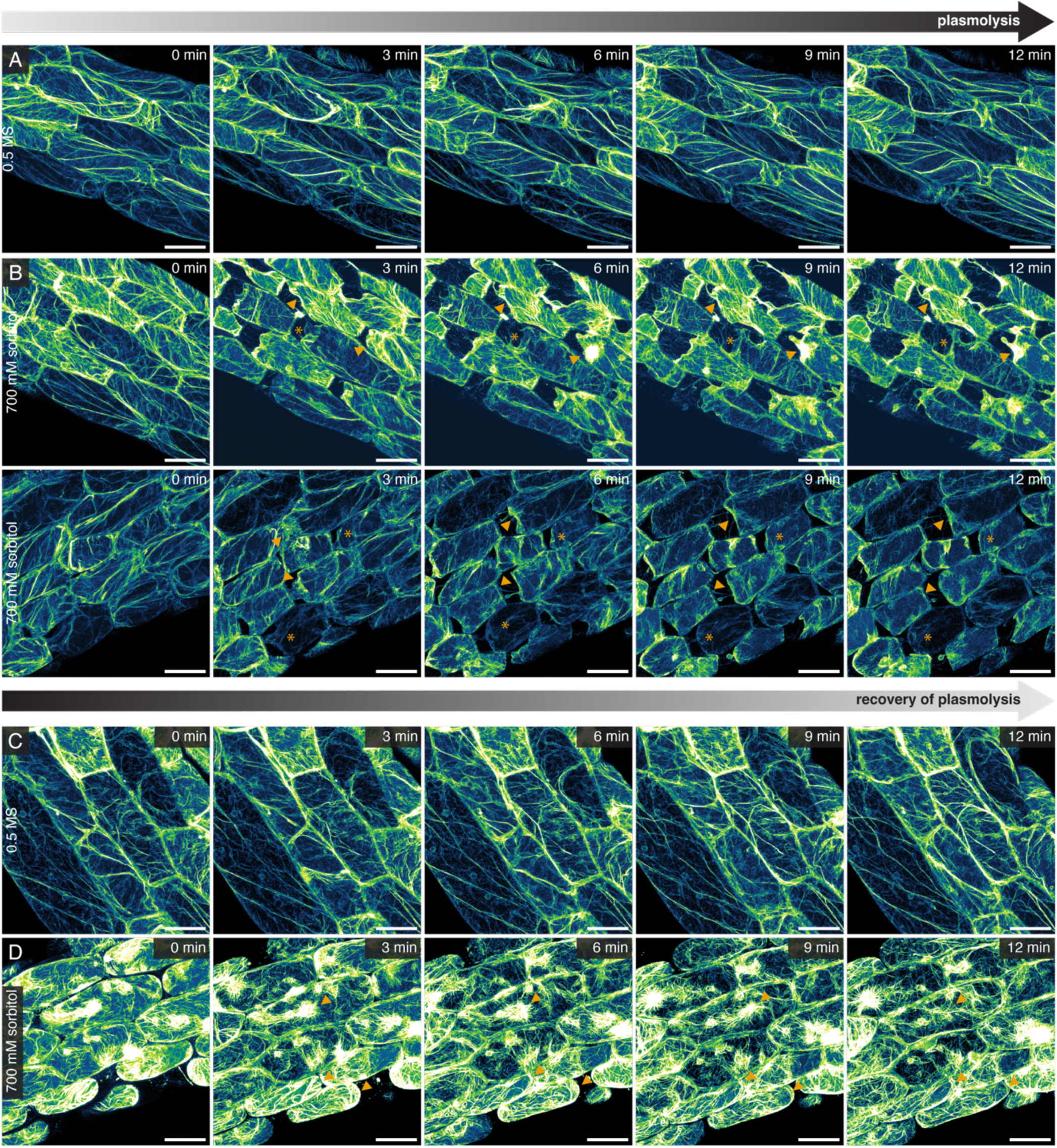
Arabidopsis actin reorganization relative to cellular deformation. (A-B) Maximum projections of series of 3 ABD2-mCherry seedlings mounted in 0.5 MS (A) or 700 mM sorbitol (B). Arrows highlight deformation sites where actin intensity increases, asterisks highlight deformation sites with no or mild increase in actin intensity. (C-D) Maximum projections of a series of 2 ABD2-mCherry seedlings that were first treated with 0.5 MS (C) or 700 mM sorbitol (D) and then mounted in 350 mM sorbitol to recover from plasmolysis. Arrows indicate sites of deformation where actin bundling and intensity decreases. Additional data is shown in Fig. S7-8. Scalebar = 25 µm.

Interestingly, for *Marchantia*, as described above, outward deformations (membrane/cell wall stretching) lead to a loss of actin, whereas here we find that as the membrane is compressed by plasmolysis, actin accumulates. We note that this is not likely merely the result of actin being compressed as a sponge by the shrinking cell, because the actin structure also visibly changes throughout this process to an increasingly bundled state. If the accumulation we observe would be the result of simple shrinkage-induced compression, we would also expect the network to adopt its original configuration after the plasmolysis is recovered; this is not the case in our experiments. Moreover, actin accumulation at inwardly deformed sites in the plant cell is also reported for very different types of deformations, including during indentation (*22*, *23*). This suggests a directionality in the mechanical awareness of the actin networks. However, since these observations are made in different models, they cannot be directly compared. Indentation (inward deformation) of hypocotyl cells in *Arabidopsis* was previously reported to lead to the local accumulation of actin (*23*), consistent with this picture. To confirm that there is indeed a directional sensing of the cellular deformations, we treated *Arabidopsis* seedlings first with either 0.5 MS (control) or 700 mM sorbitol and then mounted them in 350 mM sorbitol to recover from plasmolysis. Again, in our control treatment, actin seems to be dynamic albeit slightly more stable due to the change to 350 mM sorbitol but no apparent enrichments are visible (Fig. 6C). In the plasmolysed samples, actin is initially severely bundled (Fig. 6D). As these samples were subjected to a high osmotic stress for 30 minutes, these effects are particularly strong. We also note that in many cells, actin appears to accumulate as a dense, contracted cytoskeleton around the nucleus (Fig. 6D), to which actin is mechanically anchored (*38*, *39*). As the cells re-inflate and recover from plasmolysis by being placed in a now hypotonic solution, the observed strong bundling and enrichment at the deformation site decrease gradually to restore a native actin organisation over time (Fig. 6D). This strongly suggest that mechanically-triggered actin reorganizations are not only reversible but also sensitive to the tensorial nature of mechanical deformations.

## Discussion

We have shown that plant actin networks exhibit distinct mechano-adaptivity, changing their structure, spatial distribution, and dynamics in response to mechanical cues acting on the cell. The changes in spatial actin distribution, or, in other words, mechanically instructed actin cell polarity, were found to be an intrinsic physical property of actin networks, independent of microtubules and largely reproducible in a physical simulation model that lacks biological feedback. We found that plant actin networks are mostly responsive to large cellular deformations. This implies that actin mechano-adaptivity may be a conserved mechanical feature of eukaryotic actin networks.

Our results paint a picture of a two-geared cytoskeletal mechano-response system in plants. The microtubule array responds to subtle changes in the tissue’s mechanical field. Microtubules can sense small and slow deformations (low strain rates), and reorganize over timescales of hours in response (*19*). This response system is ideally suited for developmental mechanosensing, which relies on subtle cues that do not require immediate (rapid) intervention. By contrast, we have shown here that the mechanical responses of actin networks come into play at much larger deformations and deformation rates; and that the response in the actin network is much more rapid, occurring in tens of seconds to a few minutes. This is likely more relevant in scenarios where large deformations require rapid action, for example, to protect cells from mechanical damage during pathogenic attack (*22–25*), wounding (*40*), or osmotic stresses during drought or salt exposure (*41*). Together, these two cytoskeletal response systems enable plants to respond adequately to a range of mechanical challenges. Given the division- orientation defects in actin mutants (*42*), there may also be a yet unclear role for actin mechanosensing in subtler developmental contexts.

While we have shown that a minimal physical model can recapitulate mechano-responsive actin reorganizations during large cellular deformations, it is very likely that active biological regulation with rapid biochemical response systems is involved. In animal cells, a variety of actin-associated proteins can rapidly effectuate mechanosensing responses, for example, by the force-activated binding of so-called catch bond proteins, such as integrin (*43*), talin (*44*), vinculin (*45*), and actinins (*8*, *11*) or the mechanical activation of phosphorylation cascades, by mechanosensitive kinases such as integrin-linked kinase (*32*) or titin kinase (*46*). However, none of the known factors in the biochemical mechanoregulation of actin-binding proteins in animals have direct homologs in plants. It is thus likely that, given the vastly different cellular mechanics of plant cells and the importance of the cell wall, the green lineage has evolved independent strategies for the rapid mechano-regulation of actin structures and dynamics. Identifying these and exploring their function remains an open-ended quest.

## Materials & Methods

### Plant growth conditions

*Marchantia* and *Physcomitrium* were aseptically grown from gemmae on ½ strength Gamborg’s B5 medium (Duchefa) complemented with 0.05% MES and 1% Daishin Agar (Duchefa) and pH was adjusted to 5.7 with KOH. *Arabidopsis* (ecotype Col-0) seeds of actin marker line UBQ10::ABD2-mCherry (*47*) and 35S:LTI6b-GFP were wet sterilized with 70% ethanol for 3 minutes, rinsed with 96% ethanol and dried in sterile conditions for at least 1 hour. Seeds were sown on ½ MS medium (Duchefa) containing 0.1% 2-Morpholinoethanesulfonic acid monohydrate (MES) buffer (Duchefa), and 0.8% Plant agar (Duchefa), and the pH was adjusted to 5.7 with KOH. Seeds were stratified at 4 °C in the dark for 2 d. All plant species were grown under long-day growth conditions (16 h light, 8 h dark).

### Plasmid construction and transformations

Oligonucleotides and plasmids used and generated in this study can be found in Tables S1-2. The entry vector was prepared by PCR amplifying the pEN207 backbone using primers 077 and 078, followed by FastDigest DpnI (Thermo Fisher) restriction digestion and purification with the GeneJET purification kit (Thermo Fisher). A *Marchantia* ADF (Mp3g20570.1) was amplified from cDNA using primers 214 and 215 and, together with a PCR-amplified mScarlet3 (Addgene #189767, primer 9 and 113), placed in the linearized pEN207 backbone using HiFi Assembly (E2621, New England Biolabs). As the *Marchantia* ADFs cluster together and are thus all equally related to the same *Arabidopsis* homologs, we renamed these genes based on their location in the genome. Mp3g20570 was named ADF3, see Fig. S9). Similarly, mNeongreen was PCR-amplified using primers with overhangs (195 & 196) and inserted into a PCR linearized pEN207 backbone (primers 54 & 77) containing the LifeAct sequence via HiFi Assembly. Both entry vectors were transferred to the pGWB101 destination vector (*48*) containing the EF1α promoter via an LR Clonase II (Invitrogen) reaction. *Marchantia* was transformed using an Agrobacterium tumefaciens (GV1301) protocol as described previously (*49*). Plants were selected for positive transformations on 0.5 Gamborg B5 (1% agar) with 10 mg/L hygromycin and 100 μg/ml cefotaxime. It should be noted that the *EF1α::ADF3-mNeongreen* has been silenced over time and could not be recovered.

### Laser ablations

*Marchantia* ablations were performed on a Leica SP8-DIVE multiphoton system with a pulsed Coherent Chameleon Ti:sapphire infrared laser with a photo-stimulation module. Plants were imaged with 0.5% 488 nm (0.5%, mNeongreen) or 1% 552 nm (mScarlet3) diode laser using a ×63 (NA 1.2) objective. As the EF1α promoter showed exceptionally high expression in the rhizoid precursor cells, we ablated an adjacent cell with a single ablation point (920 nm, 350 ms, 17-22% laser power) in the middle (in the xy-plane) of the cell. The laser power was kept to a minimum to prevent drift in the z-direction and varied slightly on a day-to-day basis. For the *EF1α::LifeAct-mNeongreen* line, we recorded 100 images before the ablation and 1000 after. Images were recorded with a 0.52 seconds interval (1000 lines/second on a 512×512 image). The *EF1α::ADF3-mScarlet3* line was dim and and required more laserpower (2%) to be detectable. To prevent photobleaching we imaged this with 1 second intervals, with 20 images pre- and 500 post-bleaching. *Physcomitrium* and *Arabidopsis* were imaged as above, but using a 40x (NA 1.1) objective.

### Mechanical stimulation of Arabidopsis

*Arabidopsis* 8-day-old seedlings were subjected to either osmotic treatment or compression to mechanically stimulate them. To study actin dynamics without compression, spacers were prepared by laser cutting (Glowforge, settings speed: 210, precision power: 50, 1 pass) Hi-Bond transparent double-sided tape (RS-Pro) into little windows where the outer dimensions were 22 x 23 mm and the inner dimensions of the window were 17 x 24 mm. The thickness of the tape was 0.23 mm and two spacers were placed on top of each other to avoid any compression. Seedlings were placed in the cut window in ½ MS medium with 0.1% plant agar (Duchefa) to minimize drift of the seedling. Squeezed samples were prepared with the same medium with agar but without spacers.

Osmotic treatment was performed with either 350 or 700 mM sorbitol (Sigma) in liquid ½ MS medium (with 0.1% MES, pH 5.7). Liquid ½ MS medium (with 0.1% MES. pH 5.7) was used as a control treatment. To study dynamics in plasmolyzed tissue, seedlings were first subjected to the treatment or control treatment for 30 min, after which they were mounted in the same treatment medium without a spacer. Imaging settings are described above.

To make timeseries of actin reorganization during plasmolysis, seedlings were directly mounted in 700 mM sorbitol in liquid ½ MS medium (with 0.1% MES, pH 5.7) or liquid ½ MS medium (with 0.1% MES. pH 5.7) as control. They were then imaged for 21 minutes every 3 minutes on a Nikon C2 inverted confocal laser scanning microscope, with a 7% 561 laser (mCherry) and a ×100 (NA 1.35) objective. For the recovery of plasmolysis, seedlings were incubated in 700 mM sorbitol in liquid ½ MS medium (with 0.1% MES, pH 5.7) or liquid ½ MS medium (with 0.1% MES. pH 5.7) as control, for 30 minutes, then mounted into 350 mM sorbitol in liquid ½ MS medium (with 0.1% MES, pH 5.7). These samples were recorded with the same intervals and settings as described above. Maximum projections were made and histogram matching to correct for bleaching was performed for each timestep in ImageJ version 1.52i.

### Image analysis

Deformation and fluorescent intensity ratios were calculated with homemade Python (3.12.2) scripts using Matplotlib (3.10.0), NumPy (1.26.4), OpenCV (4.12.0), Pandas (2.2.3), PyStackReg (0.2.8), Scikitimage (0.25.0) and Scipy (1.15.1) (*50–55*). First, images were aligned using StackReg with a translation transformation. Next, we created a global mask, which allowed more robust segmentation of the individual timeframes. For this mask, all images in the timeseries were averaged, followed by thresholding using a local Otsu threshold (15 pixel radius). This binary images was used for contour finding. This filled contour was used as a mask for all images in the timeseries.

For the analysis of the timeseries, we first averaged the timeseries in blocks of 10 images, creating 110 averaged images out of 1100 images. These averaged images were masked with the global mask defined above. Next, we thresholded the images with local Otsu (15 pixels radius) and used this binary mask for contour finding. Contours were closed with 5 iterations of dilation, followed by 5 iterations of closing using a 9×9 kernel. Some timeseries required small adaptations as contours were not completely closed or merged with adjacent cells. Therefore, we wrote small case-by-case alterations, like using global Otsu thresholding, or lowering the erosion and dilation iterations. Next, we grouped the contours over time, so that e.g. “cnt#1” always corresponded to the same cell at different timepoints.

Next, we used these groups to calculate the deformation and fluorescent intensity of the cortical region (Fig 2). This works as follows; for group (e.g cnt#1), we used the contour of the last timepoint and calculated its center. We drew a line from a pixel on the contour edge to the center. From this line we extracted the first pixels in the contour (seen from the outside inwards) for a given length (1 µm as default) and averaged the pixel intensity. This allows this for the extraction of the cortical region and the average intensity. Next, we iterated over all timepoints and repeatedly selected the first pixels (with a given length) of each contour. In deformed area’s the selected pixels had different coordinates that at the start, which allowed to calculate the deformation. The intensity ratio was calculated by dividing the average intensity at a given timepoint by the average intensity of the start - 1. This whole process then iterated for all pixels at the contour edge, creating a deformation and fluorescent intensity map for every cortical pixel of every cel in the image. Signal intensity rations were grouped on their final deformation (1 µm intervals). Statistics were calculated by comparing every group to the 0 µm deformation group. Within such comparison, every timepoint was compared to the corresponding timepoint of the 0 µm deformation using pairwise t-tests with unequal variance, Multiple testing was corrected using Bonferroni’s method.

### Mathematical model

The mathematical model is described and illustrated in detail in the Supporting Information, Figures S10-12.

### Actin dynamics

Actin dynamics are quantified from confocal microscopy time-lapse sequences using an image correlation approach. Images are recorded on an inverted Leica DM8 CS Stellaris microscope equipped with a ×86 (NA 1.2) objective, with 5% 561 nm (mCherry) white light laser at a resolution of 256×256 pixels (60.31 x 60.31 microns) and a frame rate of 1.3 fps. Image correlation is performed using a home-written code in Matlab 2025b. The approach is based on the image correlation methods used in Laser Speckle Imaging (*56*, *57*). In essence, the analysis quantifies how different two image pairs, separated by a time interval Δt, are by computing the extent of image decorrelation. At short time intervals, where actin has moved little, two images are very similar and the decorrelation value is close to zero. At larger time intervals, where actin has had more time to move, decorrelation increases. This is expressed in the intensity structure function as:

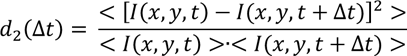

In which *I*(*x*, *y*, *t*) is a 2-dimensional image intensity matrix at time t, and the angular brackets indicate an average over all pixels in the image and over all matching image pairs in the sequence, separated by an interval Δ*t*. As these image sequences were recorded at a relatively low signal-to-noise ratio, to reduce photobleaching, images were first subjected to filtering to correct for residual photobleaching, using a histogram matching method, and to reduce spurious and fast-fluctuating cytosolic background noise, subjected to both a spatial Gaussian filter (width = 13 pixels) and a temporal rolling median filter (window size = 5 frames). For each image series, a single *_d_*_2_(Δ*_t_*) was computed; these were averaged across all replicate experiments and presented as the mean over these replicates and the standard deviation across independent replicates.

## Supporting information

Statistical data + all sequences

## Acknowledgements

We begin by expressing our gratitude to the late dr. Hanne van der Kooij for providing the inspiration to start this project. We gratefully acknowledge prof. Liam Dolan (Gregor Mendel Institute Vienna, Austria) for sharing transgenic *Marchantia* membrane and microtubule marker lines. This work and J.L., A.P., J.T., J.v.d.G., and J.S. are supported by the Dutch Research Council (NWO) through the Gravitation program GreenTE (project number: 024.006.001). J.L. and J.S. are funded by the European Research Council, project Catch (project number: 101000981).

## Author contributions

Conceptualization: J.L. / A.D. / A.P. / J.T. / U.S. / J.vd.G. / J.S.

Data curation: J.L. / A.D. / A.P./ S.G.

Formal Analysis: J.L. / A.D. / A.P.

Funding acquisition: J.vd.G. / J.S.

Investigation: J.L. / A.D. / A.P. / S.G.

Methodology: J.L. / A.D. / A.P. / J.T. / S.G. / J.S.

Project administration: J.vd.G. / J.S.

Resources: J.L. / A.D.

Software: J.L. / A.D. / A.P. / J.T. / J.S.

Supervision: J.T. / C.B. / U.S. / J.vd.G. / J.S.

Validation: J.L. / A.D. / A.P.

Visualization: J.L. / A.D. / A.P. / J.T. / J.S.

Writing-original draft: J.L. / A.D. / A.P. / J.T. / J.vd.G. / J.S.

Writing-review & editing: J.L. / A.D. / A.P. / J.T. / S.G. / C.B. / U.S. / J.vd.G. / J.S.

## Data availability

All data associated with this manuscript will be made available upon request. Data analysis codes developed the context of this work are available at: 10.5281/zenodo.21362415.

## Competing interest

We declare no competing interests

## Notes

### Competing Interest Statement

The authors have declared no competing interest.

