## Supplementary Information for "Plant actin networks are rapid mechano-adaptive scaffolds"

**This PDF file includes:**

**Supplementary Figures S1-S8**
**ADF alignment and tree construction, including Figure S9**
**Detailed description of the mathematical model, including Supplementary Figures S10-S12**

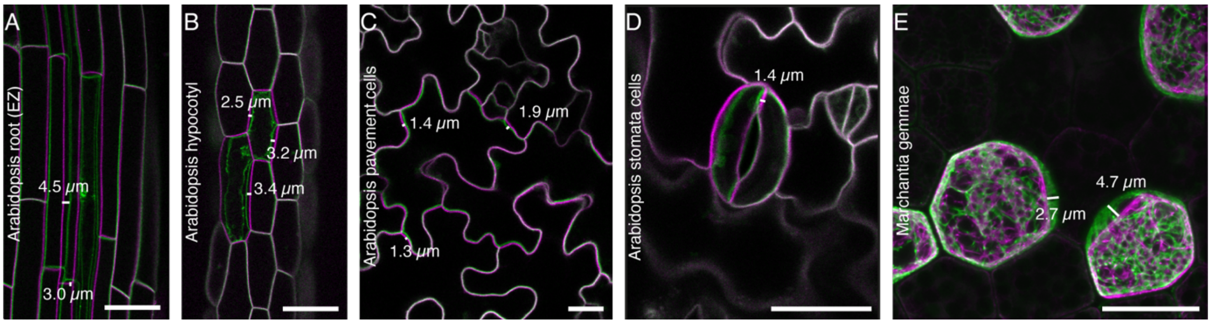


***Figure S1. The amplitude of laser ablation-induced deformations is cell-type dependent.*** *(A-E) The deformation measurements of the Figure 1. Scalebar = 25 μm.*

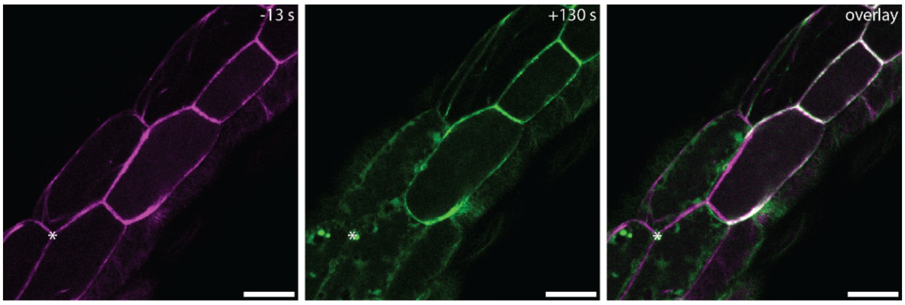


***Figure S2. Laser ablation of hypocotyl cells of ABD2-mCherry.*** *Site of laser ablation is indicated with a white asterisk. Scalebar = 25 μm.*

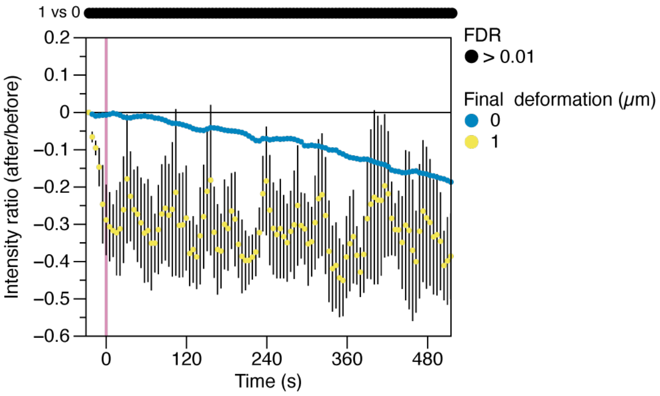


***Figure S3. The automated image analysis pipeline is validated on a no-ablation control.***

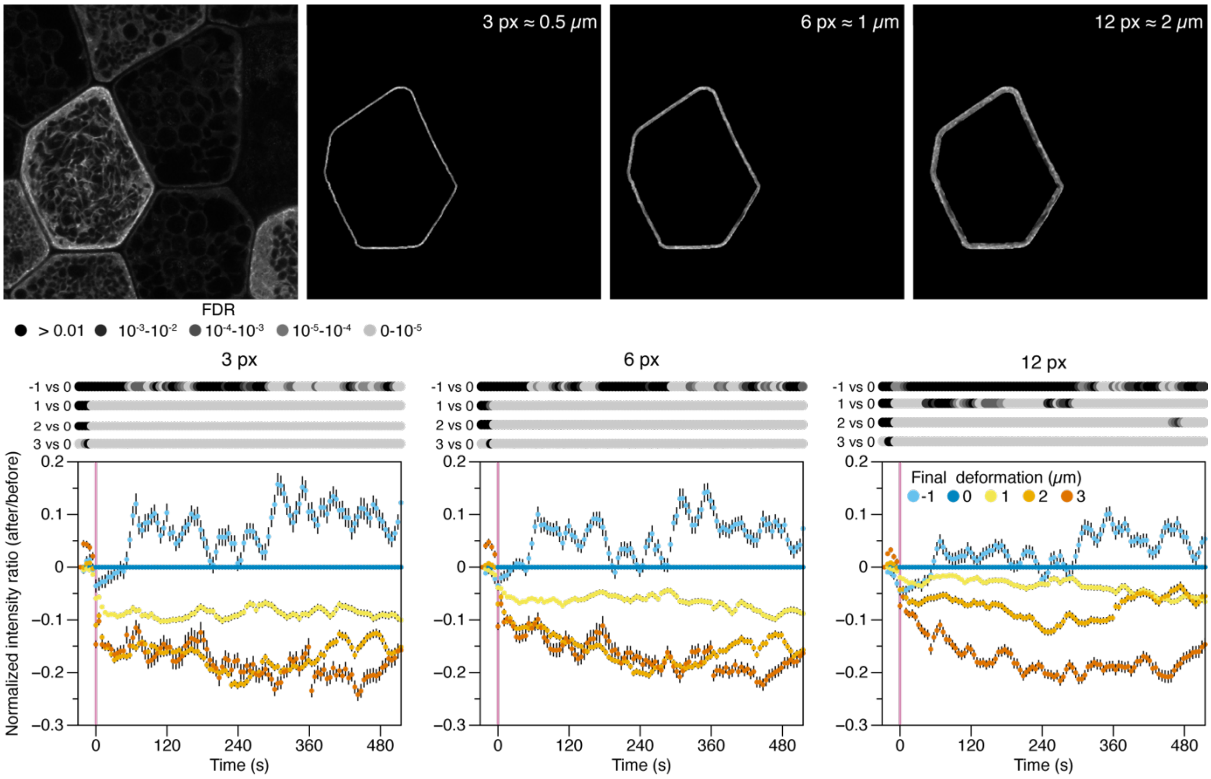


***Figure S4. The relationship between fluorescence and deformation is not dependent on the width of the cortical mask.*** *The script was tested for 3, 6 and 12 pixel wide cortical masks, but the observation that the fluorescent intensity decreased in the most deformed area's was not affected. Statistics were calculated with pairwise t tests per timepoint for every bin compared to 0 μm deformation. Multiple testing was corrected with Bonferroni's method.*

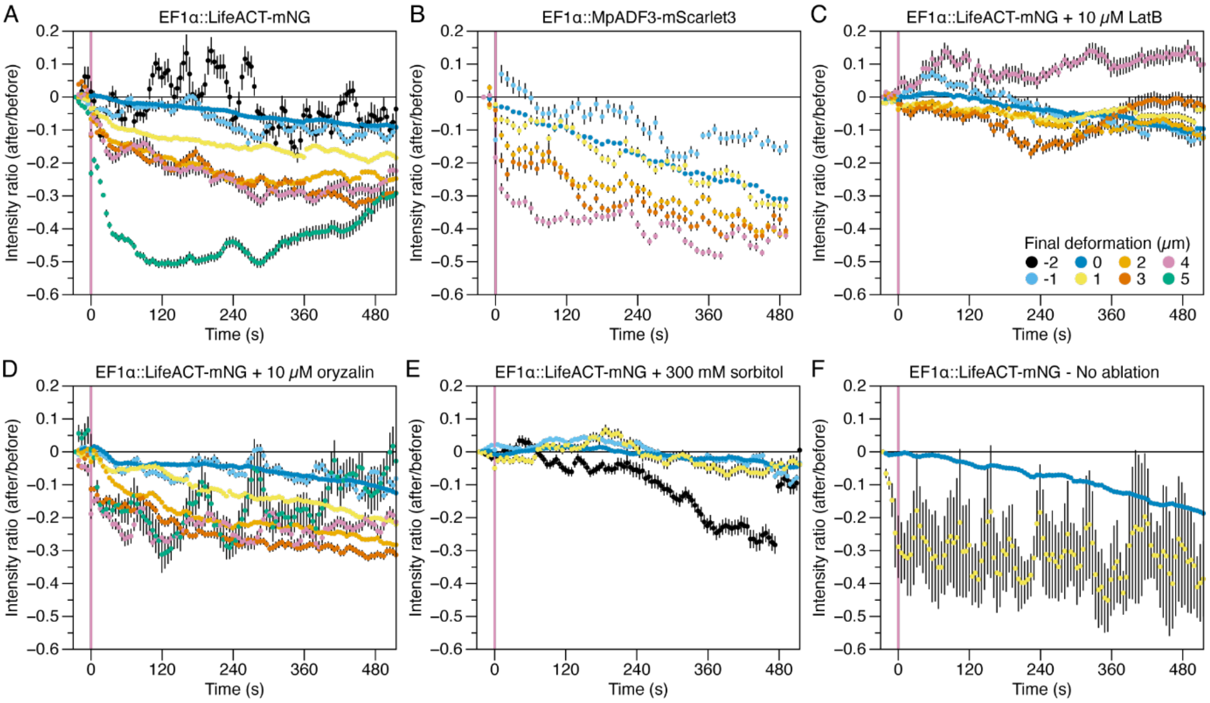


***Figure S5. Marchantia actin responds to the mechanical environment.*** *The non-normalized data as shown in Figure 2. Data was binned on the final deformation. In contrast to figure 2, all datapoints are shown with more than 50 pixels per timepoint.*

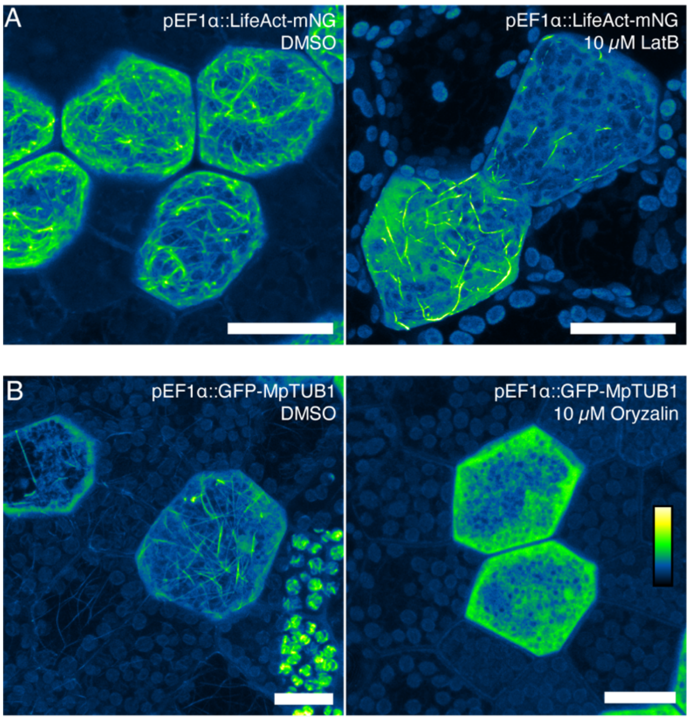


***Figure S6. The chemical treatments are functional.*** *(A) Representative images of Marchantia gemmae expressing pEF1α::GFP-MpTUB1 mounted in DMSO or 10 μM Oryzalin (A) or pEF1α::LifeAct-mNeongreen mounted in DMSO or 10 μM LatB. Scalebar = 25 μm.*

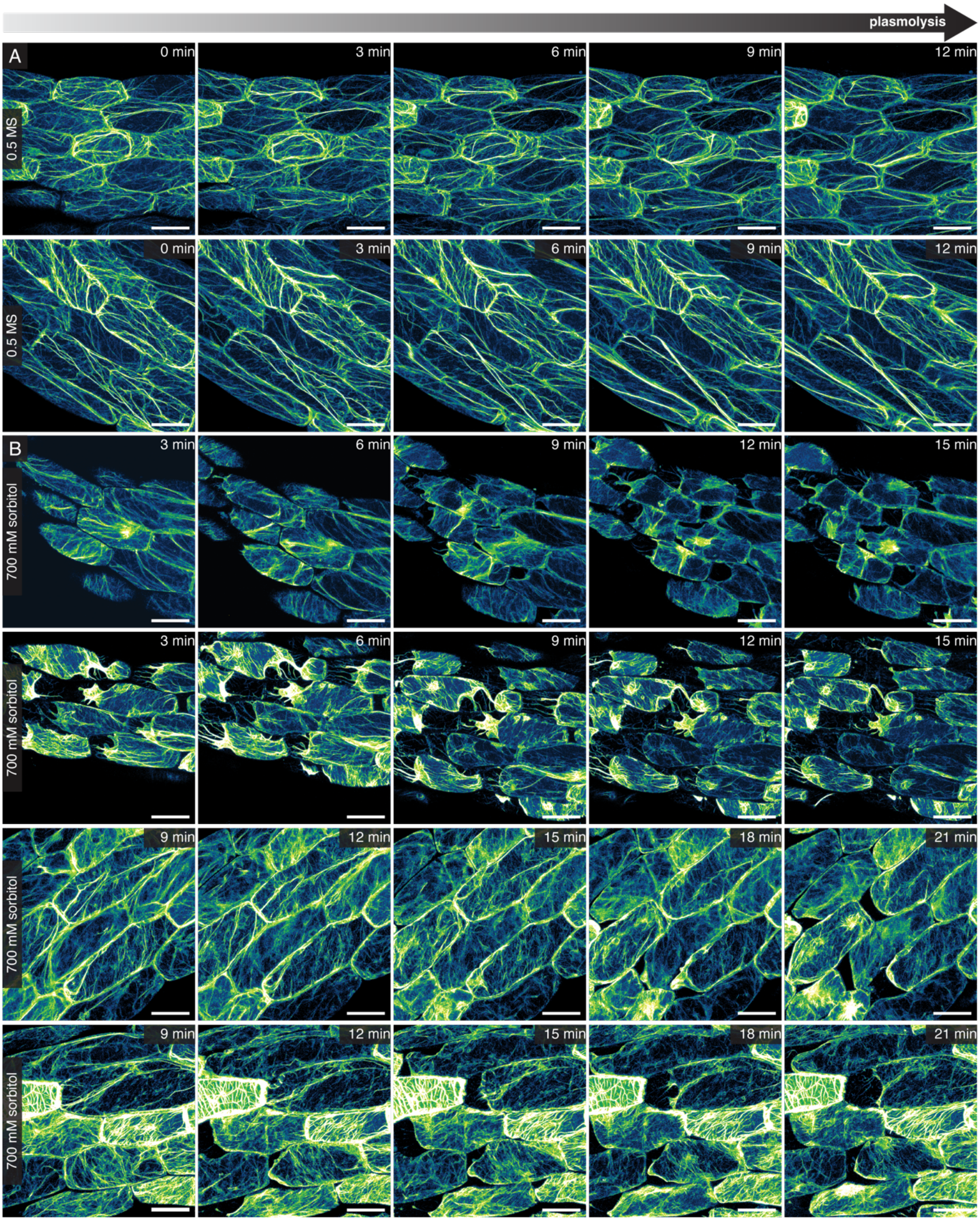


***Figure S7. Additional series of Arabidopsis actin reorganization relative to cellular deformation during plasmolysis****. Maximum projections of series of seedlings mounted in 0.5 MS (A) or 700 mM sorbitol (B). These series combined with the series shown in Fig. 6A-B are representative of the complete data set (0.5 MS n = 9, 700 mM sorbitol n = 8). Scalebars = 25 µm.*

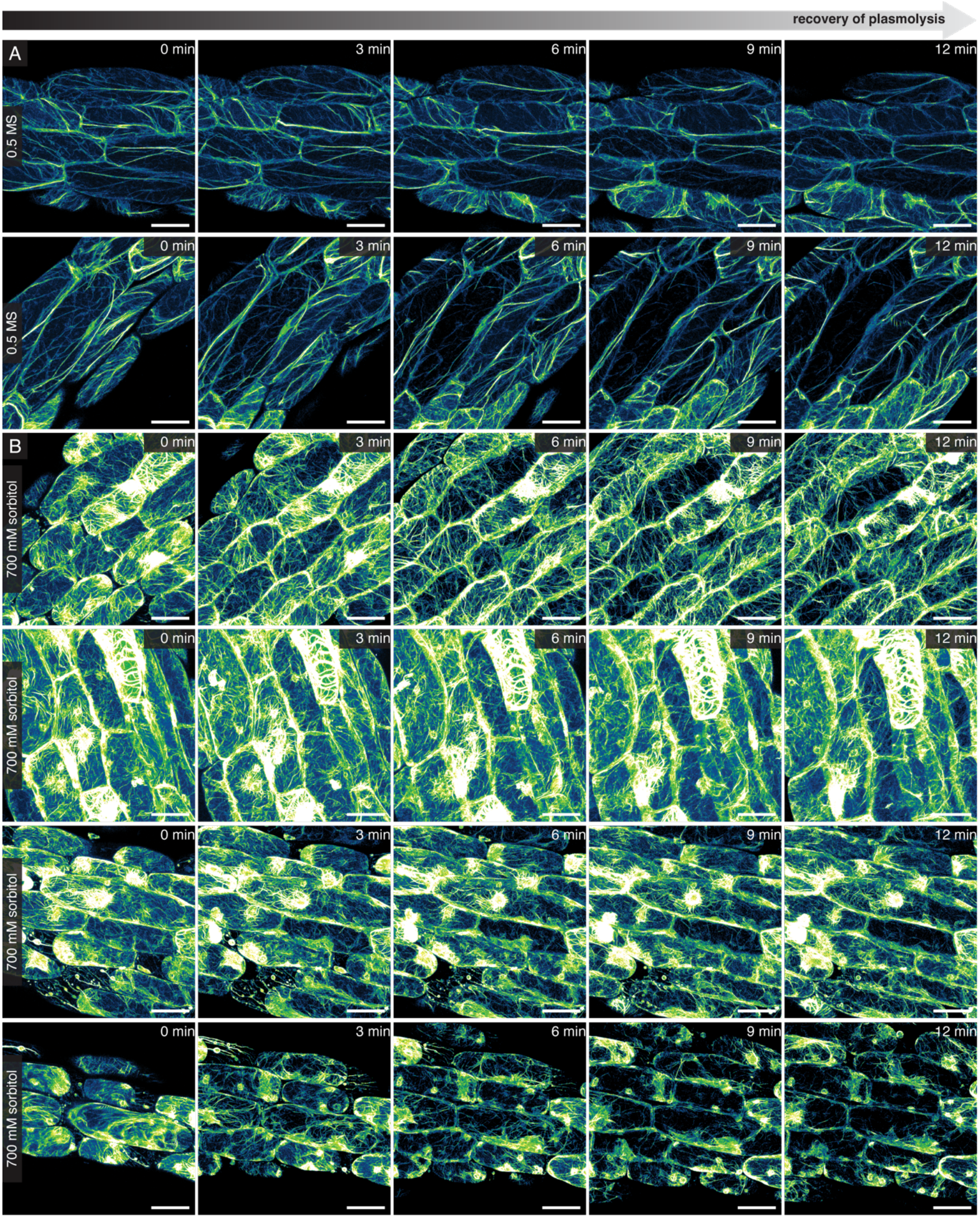


***Figure S8. Additional series of Arabidopsis actin reorganization relative to cellular deformation during recovery of plasmolysis.*** *Maximum projections of series of seedlings pre-treated in 0.5 MS (A) or 700 mM sorbitol (B) and then mounted in 350 mM sorbitol to recover from plasmolysis. These series combined with the series shown in Fig. 6C-D are representative of the complete data set (0.5 MS n = 5, 700 mM sorbitol n = 5). Scalebars = 25 µm.*

### ADF alignment and tree construction

Representative Arabidopsis ADF protein sequences were downloaded from TAIR (Arabidopsis.org) and

Marchantia sequences were identified with BLAST on the MarpolBase (Marchantia.info). Sequences were aligned with ClustalOmega on uniprot.org. The phylogenetic tree was created with IQ-TREE2 (v2.2.0)(*55*) with ModelFinder and 1000 bootstraps.

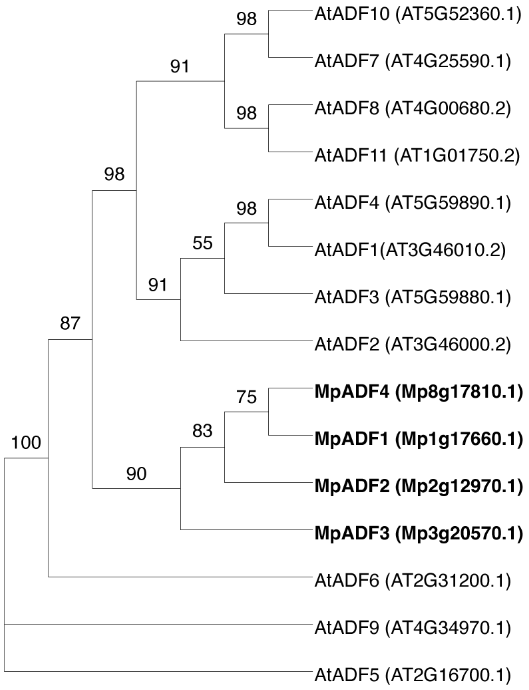


***Figure S9. Marchantia ADF proteins cluster together and are all equally related to their closest Arabidopsis homolog.*** *Phylogenetic tree of Arabidopsis and Marchantia ADF protein sequences. Numbers indicate bootstrap support.*

**Detailed description of the mathematical model**

The laser ablation experiments demonstrate how the density of the actin network responds to deflection of the cell boundary. However, they do not give access to the stresses and strains within the cytoskeleton. Hence, to predict the mechanical stress patterns that develop during the laser ablation experiments, and how they feed back into the spatiotemporal evolution of the actin cytoskeleton, we constructed a plane-strain continuum model (*1*) of the plant cell using the Finite Element Method (FEM).

**Reconstructing cell geometry from microscopy images**

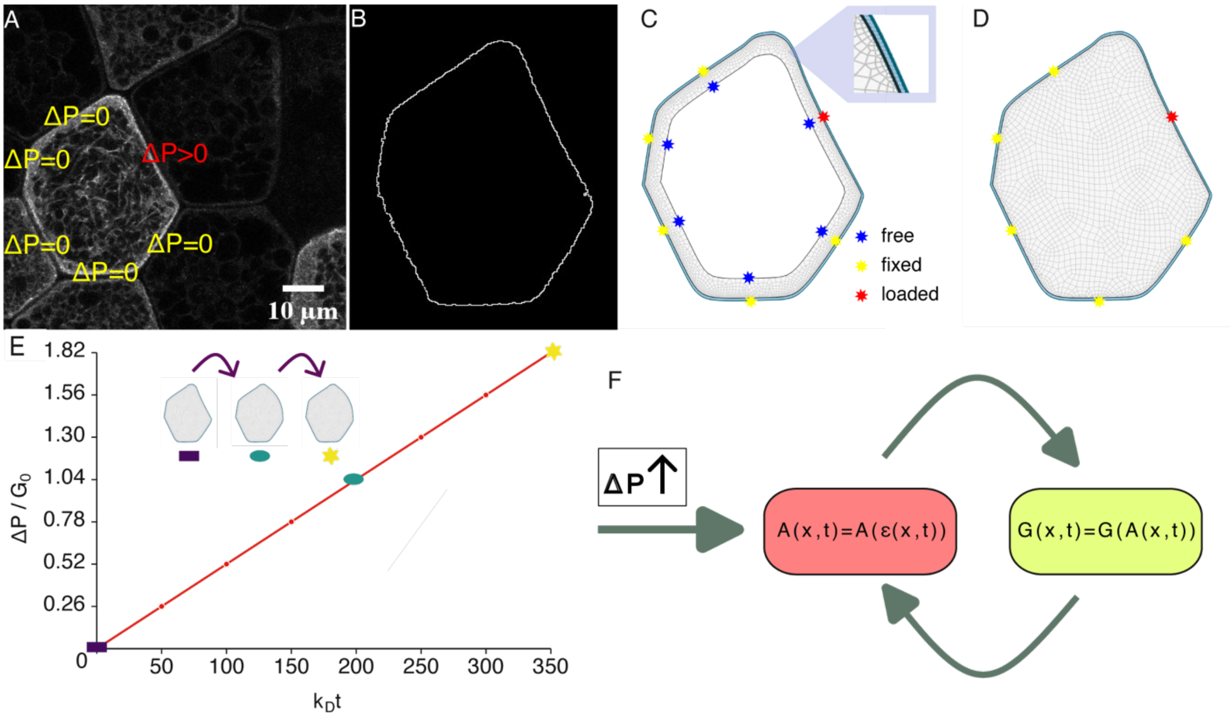


***Figure S10. Framework for modeling cell deformation from experimental observations:*** *(A) Microscopy image of the reference cell used for mesh construction. Here we show the pressure differences created at each edge of the reference cell post-laser ablation. In the simulation we gradually increase the pressure difference on the deforming cell edge to mimic experimental conditions while keeping the pressure differences on the other edges constant. (B) Cell boundary extracted through image processing in* Fiji*. (C) Shell mesh generated from the cell boundary. The enlarged view shows the blue mesh representing the cell wall and the rest of the mesh representing the cytoskeleton. The colored markers indicate the boundary conditions applied to the edges of the mesh. (D) Whole-cell mesh generated from the cell boundary. (E) Pressure difference over the loaded edge as a function of time scaled with respect to* $G_0$ *and* $k_D$*, respectively. The inset shows the shape change at three different time points. (F) Schematic of the feedback loop between the shear modulus* $G(\boldsymbol{x}, t)$ *of the cytoskeleton and the actin network density* $A(\boldsymbol{x}, t)$*.*

The model requires the geometry of the cell in the undeformed configuration as input. For this purpose, using *Fiji* (*2*), we extracted the boundary points of the cell in the undeformed configuration (see Fig. S10B) from the experimental micrographs (Fig.S10A). The extracted points were then fitted with smooth B-spline curves in *Gmsh* (*3*) to reconstruct the cell geometry. Using the reconstructed cell geometry, we generated meshes comprising two domains representing the cytoskeleton and the cell wall (Fig. S10C). Although the actin network extends throughout the cell, its density is considerably higher near the cell wall (*4*). To investigate whether the stress response and feedback depend solely on the actin cortex or also on the actin in the cell interior, we considered two limiting cases. In the whole-cell model (Fig.S10D), the entire cell interior is treated as an elastic gel. In the Shell model (Fig.S10C), only the cortical actin cytoskeleton is included.

### Model assumptions and equations

Plant tissues maintain mechanical equilibrium by balancing turgor pressure with cell wall tension and forces from neighboring cells (*5*, *6*). Laser ablation disrupts this equilibrium, creating a pressure difference (see Fig.S10A) causing the neighboring cell to expand. To represent the experimentally observed outward expansion of the cell edge, we increase the pressure difference at the corresponding edge of the mesh as shown in Fig.S10E. In the experiments, the undeformed cell edges are approximately fixed through connections via the middle lamella (*7*). Accordingly, we applied a fixed boundary condition to those edges in the simulations. Finally, in the Shell mesh the inner edge is modeled as a traction-free boundary, meaning it can move freely (see Fig.S10C-D).

To account for the large deformation observed in the experiments, we employed finite-strain theory, in which deformation is described by the deformation gradient $\mathbb{F} = \partial \boldsymbol{x} / \partial \boldsymbol{X}$. Here, $\boldsymbol{X}$ and $\boldsymbol{x}$ denote the positions of a point on the mesh in the undeformed and deformed configurations, respectively. The resistance to deformation of both the cytoskeleton and the cell wall was represented using a compressible, hyperelastic Neo-Hookean material model (*8*, *9*), with strain energy density given by Eq.(1). The energy density can be separated into a volumetric part accounting purely for volume changes and a deviatoric part accounting for shape changes:

$$W^* = \underbrace{\frac{G^*}{2}(\bar{I}_1 - 3)}_{\text{deviatoric contribution}} + \underbrace{\frac{K^*}{2}(J-1)^2}_{\text{volumetric contribution}} \quad . \tag{1}$$

Here, $W^* = W/G_0$ denotes the nondimensional strain-energy density, while $G^* = G/G_0$ and $K^* = K/G_0$ denote the nondimensional shear and bulk moduli, respectively. These quantities are normalized with respect to the shear modulus $G_0$ of the cytoskeleton at $t_0$. From the deformation gradient we can compute the first invariant $I_1 = \mathrm{tr}(\mathbb{F}^T\mathbb{F})$ and the determinant $J = \det(\mathbb{F})$. The first isochoric invariant, $\bar{I}_1 = J^{-2/3}I_1$, characterizes shape changes independent of volume change, while $J$ accounts for local compression and expansion of the network. The shear modulus and bulk modulus are kept constant for the cell wall, while the cytoskeleton is modeled with a time dependent shear modulus as a function of the actin network density (Eq. (4)) and a constant bulk modulus (Table 1).

Stress and strain are both local mechanical fields, but they can represent different biological responses (*10*–*13*). Stress describes how internal load is transmitted through the cortex, whereas strain describes how much the local network architecture is deformed. These quantities are directly proportional only for a homogeneous material with fixed elastic properties. In our coarse-grained model, however, the local stiffness depends on the actin network density ($A(\boldsymbol{x}, t)$) (4). As a result, regions with higher network density may carry larger stresses while undergoing relatively small deformation, whereas regions with lower network density may deform more strongly under a smaller load. A stress-dependent evolution law would therefore emphasize load-bearing regions, while a strain-dependent evolution law emphasizes regions where the network is geometrically distorted. From the experiments, we observe that the actin network density after laser ablation is sensitive to the deflection of the cell wall. Physically, this response may arise from deformation-induced weakening of the cortical network, for example through force-assisted molecular unbinding or depolymerization processes that destabilize the network (*14*–*16*). In our coarse-grained model, these microscopic processes are represented by a local actin network density field ($A(\boldsymbol{x}, t)$) which evolves according to a phenomenological Bell-type (*14*) strain-dependent kinetic law as follows:

$$\frac{\mathrm{d}A}{\mathrm{d}t} = k_A - k_D \exp\left(\frac{\varepsilon}{\varepsilon_{\mathrm{ref}}}\right) A + D_A \nabla^2 A \, . \tag{2}$$

The first contribution describes homogeneous actin network formation, with rate constant $k_A$. The second term accounts for network degradation, which in the absence of deformation is proportional to the network density $A$ and the degradation rate at zero strain $k_D$. Under deformation, the degradation rate increases in proportion to the ratio of a local scalar strain measure $\varepsilon$ and a reference strain $\varepsilon_{\mathrm{ref}}$ which sets the sensitivity to strain. We tested several choices for $\varepsilon$, including the strain trace, principal strain, and the deviatoric strain norm, which will be further discussed in the Results section. The third term describes diffusion and is governed by the diffusion coefficient $D_A$ (*17*). Without diffusion, Eq. (2) evolves independently at each spatial point. Even though the rest of the terms in Eq.(2) are smooth, they do not regularize spatial gradients in the actin density. We therefore include a small diffusion term to smooth the network density and improve numerical stability. We can nondimensionalize Eq. (2) to obtain:

$$\frac{\mathrm{d}A^*}{\mathrm{d}t^*} = 1 - \exp\left(\frac{\varepsilon}{\varepsilon_{\mathrm{ref}}}\right) A^* + D_A^* \boldsymbol{\nabla}^{*2} A^* \,, \tag{3}$$

here $A^* = k_D A/k_A$ , $t^* = k_D t$, $D_A^* = D_A/l_0^2 k_D$, $\boldsymbol{\nabla}^{*2} = l_0^2 \boldsymbol{\nabla}^2$ represent the nondimensionalized network density, time, diffusion coefficient and Laplacian respectively where $l_0$ is a characteristic experimental length scale. The choice of $k_A/k_D$ as the network density scale is motivated by the fact that it represents the steady-state network density in the absence of deformation-dependent degradation. Similarly, $1/k_D$ is used as the characteristic time scale, since it corresponds to the natural reaction time scale associated with network turnover.

Finally, the shear modulus of the cytoskeleton evolves as:

$$G^*(\boldsymbol{x}, t) = G^* \left(\beta A^*(\boldsymbol{x}, t) + (1 - \beta)\right), \tag{4}$$

where $\beta$ controls the sensitivity to actin network density. This equation creates a feedback loop between the evolution of the actin network density and the mechanics of the plant cell (Fig.S10F). An increase in strain leads to a decrease in the actin network density, hence decreasing the elasticity and reducing the stress in the next increment.

### Parameterization

The parameters used for the model are listed in Table 1. For comparison with experiment we pick $l_0 = 1\mu m$. The cell wall is stiffer than the rest of the cell and based on experimental values of stiffness for the cell wall (*5*) and actin cytoskeleton (*18*) the cell wall material constants are chosen to be about $10^3$ times larger than the cytoskeleton material constants. We set $\varepsilon_{\mathrm{ref}} = 1$. At the experimental length scale $l_0$, the effective diffusive rate is $D_A/l_0^2 = 2 \times 10^{-6}\ \mathrm{s}^{-1}$. This value is several orders of magnitude smaller than the reaction rates $k_A$ and $k_D$, indicating that the dynamics are dominated by reaction kinetics rather than diffusion. This conclusion remains valid even when the degradation rate is made strain-dependent, as in Eq. (3). In this case, the strain-dependent modulation changes the reaction rate only by a factor of order $10^{-1}$, which still leaves the reaction rates much larger than the diffusive rate. Therefore, network dynamics are not diffusion-limited.

### Numerical implementation

We use a monolithic transient time-stepping scheme to increase the pressure on the deforming edge to about $\Delta P/G_0 = 1.82$ over time, as shown in Fig.S10E. We choose this specific value to achieve a displacement comparable to the large instantaneous displacement observed in the experiment. To determine the deformation, we create a mapping of the deformed edge before and after laser ablation using the k-d tree method (*19*), we obtained a maximum displacement of approximately $5.6\ \mu\mathrm{m}$ and we round it off to $5\ \mu\mathrm{m}$ in the simulations. We use *pyoomph* (*20*) an open-source *C++* FEM solver with a Python wrapper to carry out the simulations.

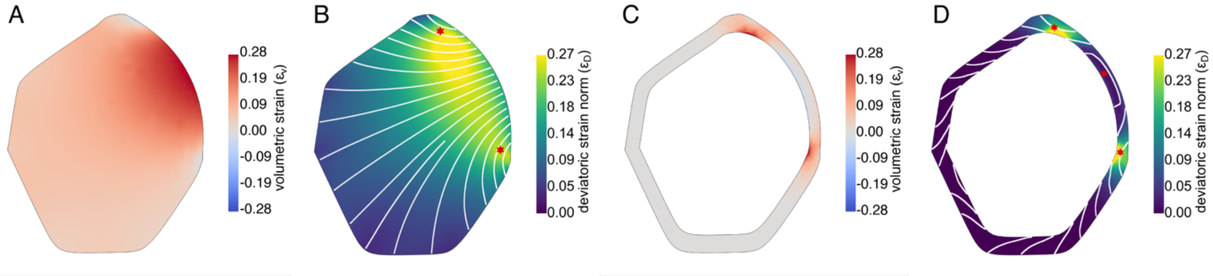


***Figure S11. Deviatoric and volumetric strain in deformed cells*** *(A) and (B) represent the volumetric and deviatoric strain, respectively, for the whole cell model, while (C) and (D) represent volumetric and deviatoric strain for the shell model.The streamlines in (B),(D) represent the principal directions which*

*represent the direction of maximum stretch. Streamlines with greater curvature are regions with higher shape distortion, as indicated by the red markers in the panels.*

| **Material Constants** | **Cell Wall** | **Cytoskeleton** |
|---|---|---|
| $G^*$ | $10^3$ | $G^*(\mathbf{x}, t)$ |
| $K^*$ | $4.33 \times 10^3$ | 4.33 |
| $\varepsilon_{ref}$ | - | 1 |
| $D_A^*$ | - | $2 \times 10^{-6}$ |
| $\beta$ | - | 0.5 |

Table 1:Nondimensionalized Material Constants used in the simulation*.*

**Deformation fields in the whole-cell versus shell model**

First, we discuss how deflection of the cell wall translates to deformation of the cytoskeleton for both the whole-cell model and the cortex model. In line with Eq. (1) we separate the deformation into contributions of shape and volumetric change. For small deformations, the deformation tensor and strain reduce to $\mathbb{F} \approx \mathbb{I} + \nabla \boldsymbol{u}, \boldsymbol{\varepsilon} = \frac{1}{2}(\nabla \boldsymbol{u} + \nabla \boldsymbol{u}^{\mathrm{T}})$, where $\boldsymbol{u}(\boldsymbol{X}) = \boldsymbol{x} - \boldsymbol{X}$ is the displacement field. Substituting this in Eq.(1) the volumetric $\varepsilon_V$ and deviatoric $\varepsilon_D$ contribution reduce to:

$$\varepsilon_V = J - 1 \approx \mathrm{tr}(\boldsymbol{\varepsilon}) = \varepsilon_{xx} + \varepsilon_{yy}. \tag{5}$$

$$\varepsilon_D = \sqrt{\frac{1}{2}(\bar{I}_1 - 3)} = \sqrt{\frac{1}{2}(I_1 J^{-2/3} - 3)} \approx \sqrt{\boldsymbol{\varepsilon}:\boldsymbol{\varepsilon} - \frac{1}{3}[\mathrm{tr}(\boldsymbol{\varepsilon})]^2}. \tag{6}$$

where $\varepsilon_{xx}$, $\varepsilon_{yy}$ are the normal strain components for x and y direction respectively, $\mathrm{tr}(\boldsymbol{\varepsilon})$ represents the trace of the strain tensor and $\boldsymbol{\varepsilon}$ represents the strain tensor. The deviatoric strain tensor is a measure of the shape distortion, excluding any volumetric contributions.

**Spatial evolution of network density**

To understand the sensitivity of actin network density to the change in volume and shape of the cell, we plot the strains given by equations Eq.(5) and Eq.(6), as shown in Fig.S11 for the whole cell mesh and the shell mesh. For the whole-cell mesh, the volumetric strain plot clearly shows expansion near the center of the cell close to the deformed edge and compression near the corners of the deformed edge (Fig. S11A). In contrast, the cortical mesh shows maximum expansion near the corners of the interior deformed boundary, while the central region of the interior deformed boundary is weakly compressed (Fig. S11C). This difference is physically intuitive: in the whole-cell mesh, the deformed boundary is connected to the central part of the cytoskeletal network, which leads to extension of the central region of the deformed edge. By contrast, this interior connection is absent in the cortical mesh, allowing the thin cortical shell to undergo comparable deformation with relatively small volumetric changes in the central region of the deformed edge.

The deviatoric strain norm in the whole-cell mesh is higher close to the corners of the deformed edge, while it remains lower along the rest of the deformed edge. (Fig. S11B). It is interesting to note that the shear is highest slightly below the deformed edge, close to the top corner of the deformed boundary. (Fig. S11B). The streamlines in the figure indicate the principal direction, i.e. the direction of maximum stretch. Principal directions with higher curvature indicate regions of larger shear and hence larger shape changes. As seen from (Fig.S11B), this confirms that shape distortion is higher near the corners of the deformed edge. Similar results are observed for the cortical deviatoric strain norm (Fig.S11D).

Although the overall deformation of the cell boundary is comparable between the two meshes, the local strain fields differ substantially. This leads to a different network density distribution, as shown in Fig.S12.

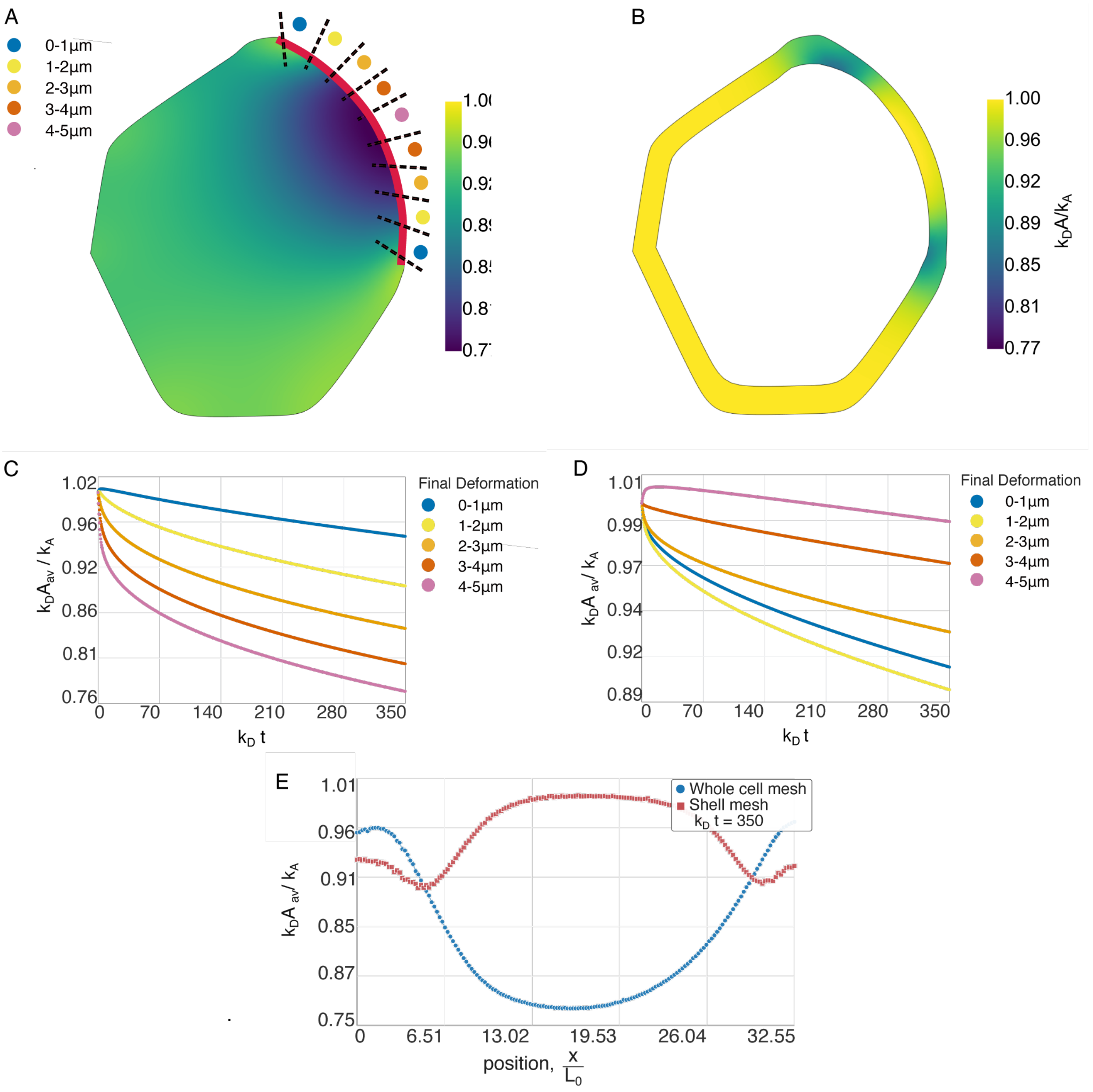


***Figure S12. Bound network density distribution and temporal evolution.*** *(A, B) Two-dimensional heatmaps of the network density for the whole-cell mesh and shell mesh, respectively. For plots(C, D, E) we evaluate the network density along the deformed edge by dividing the boundary into deformation intervals of* $1\mu m$, as indicated in (A). (C, D) Temporal evolution of the average bound network density for the whole-cell mesh and shell mesh, respectively. (E) Comparison of the network density variation along the deformed edge for the whole-cell and shell meshes at $k_D t = 350$.
